# Data-driven discovery of gene expression markers distinguishing pediatric acute lymphoblastic leukemia subtypes

**DOI:** 10.1101/2024.02.26.582026

**Authors:** Mona Nourbakhsh, Nikola Tom, Anna Schrøder Lassen, Helene Brasch Lind Petersen, Ulrik Kristoffer Stoltze, Karin Wadt, Kjeld Schmiegelow, Matteo Tiberti, Elena Papaleo

## Abstract

Acute lymphoblastic leukemia (ALL), the most common cancer in children, is overall divided into two subtypes, B-cell precursor ALL (B-ALL) and T-cell ALL (T-ALL), which have different molecular characteristics. Despite massive progress in understanding the disease trajectories of ALL, ALL remains a major cause of death in children. Thus, further research exploring the biological foundations of ALL is essential. Here, we examined the diagnostic, prognostic, and therapeutic potential of gene expression data in pediatric patients with ALL. We discovered a subset of expression markers differentiating B- and T-ALL: *CCN2*, *VPREB3*, *NDST3*, *EBF1*, RN7SKP185, RN7SKP291, SNORA73B, RN7SKP255, SNORA74A, RN7SKP48, RN7SKP80, LINC00114, a novel gene (ENSG00000227706), and 7SK. The expression level of these markers all demonstrated significant effects on survival of the patients, comparing the two subtypes. We also discovered four expression subgroups in the expression data with eight genes driving separation between two of these predicted subgroups. A subset of the 14 markers could separate B- and T-ALL in an independent cohort of patients with ALL. This study can enhance our knowledge of the transcriptomic profile of different ALL subtypes.

## 1 Introduction

Acute lymphoblastic leukemia (ALL) is a hematological cancer and the most common cancer in children, with a prevalence of ∼25% of cancers in children below 15 years of age [1,2]. ALL is diagnosed by studying cell morphology, immunophenotype, genetics/cytogenetics, and genomics and is treated with chemotherapy, targeted therapies, and antibodies [3]. ALL has a high overall survival rate, having remarkably improved from ∼10% in the 1960s to ∼90% today [4]. Reasons for this increase include optimized chemotherapy regimens, risk-based therapy, and the emergence of targeted therapies [5]. Nevertheless, ALL remains a major cause of death in children with cancer [6]. Thus, further research delving into the biological underpinnings of ALL is needed.

Based on immunophenotyping, the two major subtypes of ALL include B-cell precursor ALL (B-ALL) and T-cell ALL (T-ALL), accounting for approximately 85% and 15% of pediatric ALL cases, respectively [7]. Chromosomal aberrations and single nucleotide variants frequently occur in B-ALL. For example, hyperdiploidy, amplifications, translocations, and deletions have been observed [8,9] and single nucleotide variants and indels have been reported in transcription factors, epigenetic regulators, cell cycle regulators, and RAS pathway genes [10]. T-ALL is characterized by oncogenic NOTCH signaling due to activating mutations in *NOTCH1* [11] and abnormal expression of transcription factors due to chromosomal rearrangements [12,13]. Similarly to B-ALL, mutations and deletions have been observed in cell cycle regulators, tumor suppressors, epigenetic factors, and regulators of other signaling pathways such as JAK/STAT, PI3K, and MAPK signaling [14–19].

While immunophenotyping distinguishes the two major subtypes of ALL, several studies have further elucidated the heterogeneity and complexity within these subtypes. Multiple subgroups within B- and T-ALL have been reported based on gene expression profiling, chromosomal alterations, or DNA methylation patterns [20–25].

The molecular characteristics of B- and T-ALL have primarily been enabled by advances in next-generation sequencing technologies, particularly transcriptomics. RNA sequencing (RNA-seq) has previously been used to discover novel ALL subtypes and for diagnostic purposes [5,26–28]. Thus, understanding the information stored within ALL transcriptomics is essential. We now have access to several -omics data from pediatric cancer samples deposited in various public databases. For instance, the Therapeutically Applicable Research to Generate Effective Treatments (TARGET) project aims to identify molecular alterations driving pediatric cancers to pinpoint novel therapeutic targets and prognostic markers. This initiative has made considerable progress in our knowledge of childhood cancers (https://www.cancer.gov/ccg/research/genome-sequencing/target).

In this study, we have applied a data-driven approach to examine the diagnostic, prognostic, and therapeutic potential of gene expression data in pediatric patients with ALL. Specifically, the aim of this study is 1) to discover gene expression markers that differentiate the two major ALL subtypes, B- and T-ALL, 2) to explore the prognostic and therapeutic potential of the predicted gene expression markers, and 3) to predict further subgroups beyond these two overall subtypes. For this purpose, we have analyzed gene expression data of a pediatric ALL cohort from TARGET and validated these findings in an independent cohort of Danish pediatric patients with ALL (**Figure 1**). This study can improve our biological understanding of the transcriptomic profile of ALL subtypes. GitHub and OSF repositories associated with this study are available at https://github.com/ELELAB/ALL_markers, https://github.com/ELE-LAB/RNA_DE_pipeline, and https://osf.io/kgfpv/.

**Figure 1.**
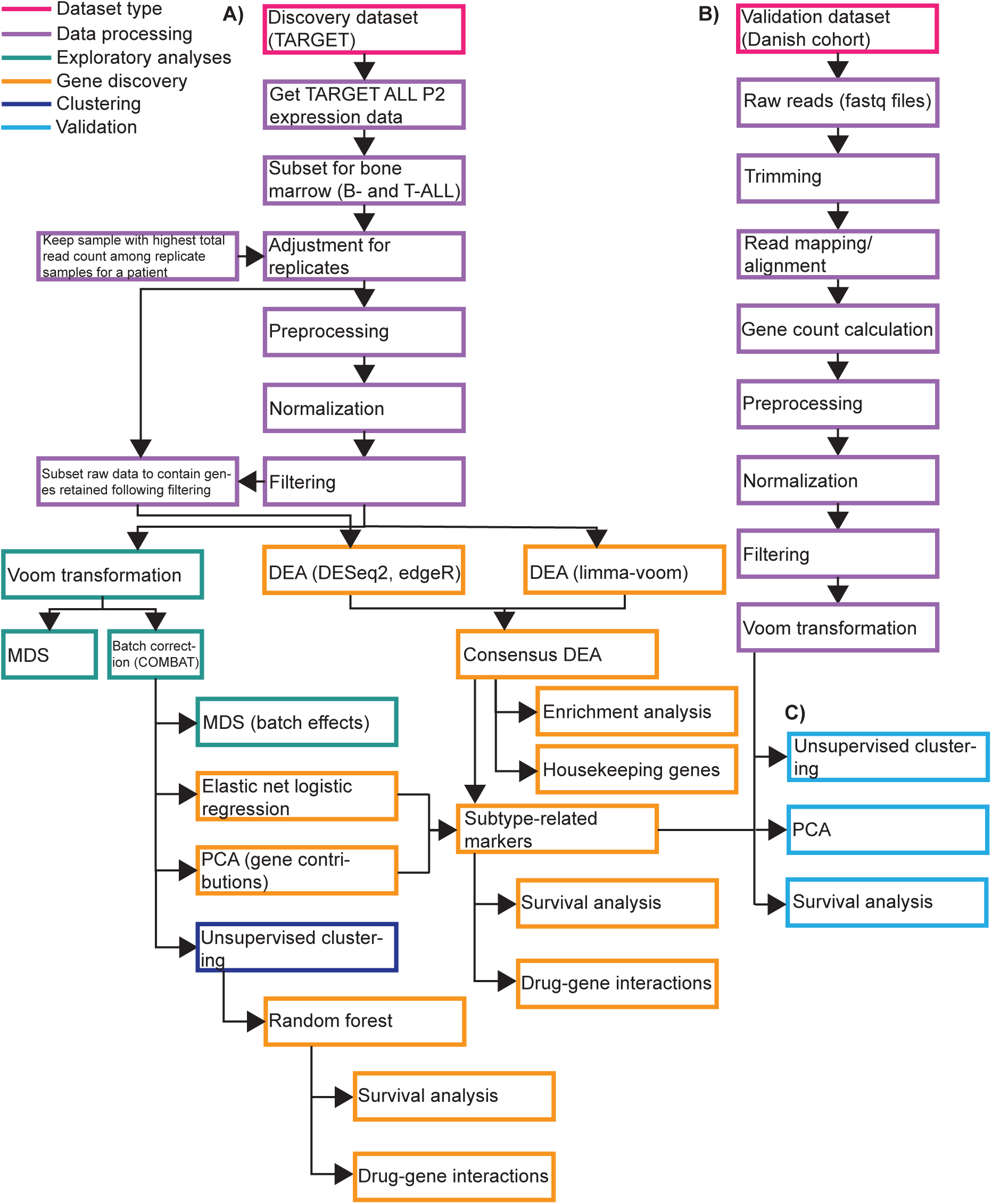
Workflow of the presented study. Each box represents an analysis which is colored according to the type of analysis. **A)** Workflow of analyses performed on the TARGET-ALL-P2 discovery dataset. **B)** Workflow of how the independent validation dataset of a Danish cohort of pediatric patients with ALL was analyzed from raw data to a gene expression matrix. **C)** Workflow of in silico validation of the predicted results from the discovery dataset in the independent cohort.

## 2 Methods

### 2.1 Download and processing of RNA-seq data of the TARGET-ALL-P2 project

We downloaded and aggregated the RNA-seq data from the TARGET-ALL-P2 project which can be accessed at National Cancer Institute’s Genomic Data Commons (http://gdc.cancer.gov) using the *GDCquery, GDCdownload*, and *GDCprepare* functions from TCGAbiolinks [29–31]. Additionally, we obtained subtype, gender, vital status, and age information using the *primary_diagnosis*, *gender*, *vital_status*, and *age_at_diagnosis* variables available in the downloaded *SummarizedExperiment* object of the data, respectively. We retained the primary samples from bone marrow for analysis only, as this was the most extensive available dataset that would ensure that the direct comparison between the two subtypes would not be confounded by differences in tissue type and recurrence. In particular, we analyzed 387 samples, of which 245 samples belonged to T-ALL and 142 to B-ALL. An overview of the samples belonging to combinations of tissue source, recurrence, subtype, and age distribution of retained samples is reported in **Supplementary Figure S1**.

In addition, we identified nine patient samples that had two replicates each. We thus retained only those with the highest total read counts to prevent bias in the downstream analyses, resulting in 378 samples (133 B-ALL and 245 T-ALL samples). More information on how the replicates have been analyzed is reported in the GitHub and OSF repositories associated with the study. Moreover, to ensure proper batch effect design, we explored the number of samples available for each annotation that can be used to describe batch factors (**Table 1**).

**Table 1.** Overview of acute lymphoblastic leukemia (ALL) samples from TARGET including information on subtypes, year of diagnosis, and tissue portion. The data refers to 133 B-cell precursor ALL (B-ALL) and 245 T-cell ALL (T-ALL) tumor samples.

|  |  | Total number of samples | Number of B-ALL samples | Number of T-ALL samples |
| --- | --- | --- | --- | --- |
| Year of diagnosis | 2004 | 10 | 10 | 0 |
|  | 2005 | 39 | 39 | 0 |
|  | 2006 | 33 | 33 | 0 |
|  | 2007 | 60 | 20 | 40 |
|  | 2008 | 47 | 18 | 29 |
|  | 2009 | 93 | 12 | 81 |
|  | 2010 | 74 | 1 | 73 |
|  | 2011 | 22 | 0 | 22 |
| Tissue portion | A | 341 | 98 | 243 |
|  | B | 37 | 35 | 2 |

Abbreviations: B-ALL, B-cell precursor ALL; T-cell ALL, T-ALL.

Next, we preprocessed the data using the *TCGAanalyze_Preprocessing* function from TCGA-biolinks [29,31]. Here, we removed outlier samples based on pairwise Spearman correlation coefficients with a cutoff of 0.6 as done in the original The Cancer Genome Atlas (TCGA) workflow [31]. We normalized the data based on GC content and library size using the *TCGAanalyze_Normalization* function from TCGAbiolinks [29,31] as these factors might bias differential expression results [32]. We used an updated version of GC content annotations (15/04/2022) as the original table in TCGAbiolinks led to the loss of too many ENSEMBL gene IDs due to a lack of annotations. The changes have been included in TCGAbiolinks version 2.24.2 (**Supplementary Text S1**, **Supplementary Table S1**). In the processing before DEA, we filtered lowly expressed ENSEMBL gene IDs using TCGAbiolinks’ function *TCGAanalyze_Filtering* as these might be artifacts or noise. Studies have reported improved sensitivity and power of DEG detection following filtering of lowly expressed genes, a step recommended before DEA, for example, with limma-voom [33–35]. In this filtering step, we used a quantile filtering with the 25th quantile as the threshold as done in the original TCGA workflow and previously used [30,31] (**Figure 1**). Following data processing, the filtered count matrix contained 42271 ENSEMBL gene IDs and 378 samples (133 B-ALL and 245 T-ALL samples).

### 2.2 Differential expression analysis (DEA) between ALL subtypes

We performed DEA between the B- and T-ALL subtypes using three different methods: DESeq2 [36], limma-voom [34,37], and edgeR [38]. We found the “tissue portion” and “year of diagnosis” variables as possible batch effects based on exploratory data analyses with the multidimensional scaling (MDS) method for dimensionality reduction (see GitHub repository). In this case, “tissue portion” refers to different portions of the original biological sample whose material was used for the RNA-seq experiments and can have value A or B. Thus, we included these as covariates in the design for DEA with the limma-voom [34,37], edgeR [38], and DESeq2 [36] methods. In more detail, the MDS analysis revealed that samples were partitioned into three main clusters, with one of them being composed entirely of B-ALL samples labeled as portion B, representing a potential batch effect, possibly introduced by technical differences in tissue collection or storage between portion A and B samples. Moreover, displaying the year of diagnosis of each sample revealed that one of two of the aforementioned clusters included B-ALL samples from patients diagnosed in the earlier years (2004-2007), while the other two clusters mainly included both B-ALL and T-ALL samples for patients diagnosed in later years (2007-2011). We also realized that the initial goal of the TARGET project was to characterize B-ALL samples alone, which was later extended to include the T-ALL subtype as well (https://gdc.cancer.gov/content/target-all-publications-summary). This variable might underlie differences in techniques and protocols performed between 2004 and 2011, representing another batch effect. Thus, we performed DEA using four different designs in limma-voom, edgeR, and DESeq2: 1) conditions (B-ALL vs T-ALL), 2) conditions and tissue portion, 3) conditions and year of diagnosis, and 4) conditions, tissue portion and year of diagnosis. In limma-voom, we transformed the data using the *voom* function from the limma package before DEA. We fitted a linear model to the expression data for each gene using the *lmFit* function, and an empirical Bayes method was used to assess differential expression using the treat approach with log2 fold change (log2FC) >= 1. In DESeq2 and edgeR, we used raw counts subsetted to contain the same ENSEMBL gene IDs as the filtered count data in limma-voom. In edgeR, DEA was carried out using the standard workflow in which a quasilikelihood negative binomial generalized log-linear model was fitted to the gene expression data using the *glmQLFit* function, and threshold testing for differential expression was performed using the treat method with log2FC >= 1. In DESeq2, differential analysis was performed with the standard DESeq2 workflow with increased iterations in the nbinomWaldTes function (maxit = 500). Threshold testing with log2FC >= 1 was specified with the results function. In all DEAs, we selected ENSEMBL gene IDs with False Discovery Rate (FDR) <= 0.05 as significantly differentially expressed. We converted ENSEMBL gene IDs into gene names using the biomaRt R package [39]. We visualized intersections between DEGs predicted by the three DEA pipelines when using four different designs using the UpSetR R package [40]. We performed a one-way ANOVA to test for statistical significance between log2FC values of consensus DEGs predicted by limma-voom, DESeq2, and edgeR.

### 2.3 RNA-seq pipeline of ALL samples from a Danish cohort

We analyzed RNA-seq data from 105 samples of ALL provided by the Rigshospitalet (Denmark) [41–43]. We designed a Snakemake pipeline [44] to obtain read counts from the raw reads of these samples. The code is available through our GitHub repository: https://github.com/ELELAB/RNA_DE_pipeline. The analyses were carried out with the pipeline version available on 1st November 2021. The workflow indexed the reference genome hg38 using STAR [45] and GENCODE transcript annotations. Raw reads were trimmed for adapters, filtered on length using Cutadapt [46], and aligned onto human reference genome hg38 using STAR. Alignments were sorted using Picard (http://broadinstitute.github.io/picard). We estimated the gene counts using FeatureCounts from the SubRead package [47]. Quality control (QC) of the input raw reads was done using FastQC [48]. QC metrics based on BAM files were provided by Picard tools, by the RSeQC package [49]. We aggregated the QC results in a single report using MultiQC [50]. More details about the pipeline settings are provided in **Supplementary Text S2**. After QC, we retained 88 samples for analyses, of which 77 and 11 belonged to the B- and T-ALL subtypes, respectively.

### 2.4 Data analysis of ALL samples from a Danish cohort

We processed the resulting gene expression data described above with preprocessing, normalization, filtering, and voom transformation. We performed unsupervised hierarchical clustering of the expression data with the complete method and Euclidean distance and visualized the results in heatmaps using the gplots R package [51]. We conducted principal component analysis (PCA) using the factoextra and FactoMineR R packages [52,53] and survival analysis using the R packages survminer [54], survival [55,56], and survMisc [57]. For survival data, we used the patients’ vital status (alive or dead) and survival time calculated as the time difference in years between 2024-01-11 and the diagnosis date for alive patients and as the time difference in years between the date of death and diagnosis date for dead patients. We applied Cox proportional hazards regression as detailed below.

### 2.5 Feature selection using elastic net logistic regression

We performed elastic net binomial logistic regression using the cv.glmnet function from the glmnet R package [58] and the approach outlined in previous work [59]. As part of the exploratory data analyses described above, we batch corrected the filtered data for the year of diagnosis variable using the function *TCGABatch_Correction* from TCGAbiolinks [30]. We used this batch-corrected data as input for elastic net logistic regression. We encoded the dichotomized target variable as 0 corresponding to B-ALL and 1 to T-ALL. We used 5-fold crossvalidation with misclassification error as the loss function and 0.5 as the elastic net mixing parameter. We used a quarter of the B-ALL samples and a quarter of the T-ALL samples as a test dataset (96 samples). The remaining samples generated the training data (282 samples). We obtained the prediction misclassification error by comparing the predictions of the trained model on the test data with the actual class labels. We performed elastic net logistic regression 10 times using 10 random seeds. We retained those ENSEMBL gene IDs selected as features in all 10 runs, thereby creating an intersected set of selected ENSEMBL gene IDs. We calculated the average elastic net coefficients of the intersected set of selected ENSEMBL gene IDs across the 10 seeds run. We converted ENSEMBL gene IDs into gene names using the biomaRt R package [39] and retrieved biotype information from the ENSEMBL database (ensembl.org).

### 2.6 Feature selection using random forest

We conducted feature selection with random forest using the R packages varSelRF and randomForest [60–62] as previously done [59] and implemented in the CAMPP2 package (https://github.com/ELELAB/CAMPP2), the second version of CAMPP published in [63]. For the feature selection process, we used 5000 decision trees for the first forest and 2000 trees for all additional trees as recommended [60,64]. At each iteration, we excluded 20% of the features from those used in the previous forest as suggested [60,64]. The least important features were excluded at each iteration. The out-of-bag (OOB) errors from all fitted random forests were explored to select the final features. The final model was selected as the one containing the least amount of features with an OOB error within one standard error of the minimum OOB error of all fitted random forests. We repeated the feature selection process 10 times using 10 random seeds and retained those ENSEMBL gene IDs selected in all 10 runs. We converted ENSEMBL gene IDs into gene names using the biomaRt R package [39] and retrieved biotype information from the ENSEMBL database (ensembl.org).

### 2.7 Feature contributions from PCA

We carried out PCA using the factoextra and FactoMineR R packages [52,53] to investigate which ENSEMBL gene IDs contributed the most to the first two principal components (PCs). Similarly to elastic net logistic regression, we used batch-corrected data as input. We investigated the top 40 ENSEMBL gene IDs contributing the most to PC1 through the *fviz_contrib* and *facto_summarize* functions. We converted ENSEMBL gene IDs of the resulting 40 ENSEMBL gene IDs into gene names using the biomaRt R package [39] and retrieved biotype information from the ENSEMBL database (ensembl.org).

### 2.8 Enrichment analyses of ENSEMBL gene IDs

We performed enrichment analyses of ENSEMBL gene IDs using the enrichR R package [65–67]. We used the following databases for the enrichment analyses: GO Molecular Function 2021, GO Biological Process 2021, and MSigDB Hallmark 2020.

### 2.9 Unsupervised consensus clustering on gene expression data using cola

We conducted unsupervised consensus clustering using the R/Bioconductor package *cola* [68] on two data inputs: 1) batch-corrected gene expression data and 2) raw gene expression data where replicates have been adjusted for. We processed the raw data using *cola’s* pre-processing function *adjust_matrix*, which imputes missing values, adjusts outliers, and removes rows with very small variance [68]. To perform the consensus clustering, we applied *cola*’s *run_all_consensus_partition_methods* function, which runs 20 different feature selection methods and partitioning combinations. The four feature selection methods used were standard deviation (SD), median absolute deviation (MAD), coefficient of variation (CV), and ability to correlate to other rows (ATC). The five partitioning methods applied were hierarchical clustering (hclust), *k*-means clustering (kmeans), spherical *k*-means clustering (skmeans), model-based clustering (mclust), and partitioning around medoids (pam). For all 20 methods, we investigated the number of clusters for *k* ranging from 2 to 6. We generated an HTML report of all results using *cola’s* function *cola_report*. We compared the performance of the 20 methods and the batch corrected and raw *cola*-processed data in three ways: 1) comparison of *k* = 2 clusters with the already annotated class labels of B- and T-ALL, 2) statistical metrics provided from the *cola* analysis: the 1-the proportion of ambiguous clustering (1-PAC) score, mean silhouette score, and concordance, and 3) visual inspection of consensus heatmaps illustrating the stability of the subgrouping provided from the *cola* analysis. After investigating these three criteria, we selected the optimal method and its optimal *k* for the final clustering of the data.

### 2.10 Survival analysis

We performed survival analysis of the gene expression markers using the R packages survminer [54], survival [55,56], and survMisc [57]. As survival data, we used the patients’ last follow-up date and days to death and vital status (alive or dead). First, we applied Cox proportional hazards regression analysis to model the effect of gene expression on survival with gene expression as a continuous independent variable and survival data as the response variable. Here, we first tested the proportional hazards assumption via the *cox.zph* function and kept only genes satisfying this assumption. These genes were afterward subject to a univariate Cox regression analysis with the *coxph* function. We corrected the *p*-values for multiple testing using the FDR method and kept those genes whose expression significantly affected survival (FDR < 0.05). Subsequently, we fit a multivariate Cox regression model on these genes, accounting for the age and sex of patients as covariates. We deemed those genes whose expression significantly affected survival from the multivariate analysis as prognostic (*p*-value < 0.05). Furthermore, we conducted a Kaplan-Meier survival analysis on the prognostic genes to assess variations in survival between two distinct expression groups. Patients were categorized into high and low-expression groups based on whether their expression values were above or below the median expression level of the corresponding gene. Survival curves were constructed using the discrete expression group as the independent variable, and the significance of the difference in survival between the two groups was assessed using a log-rank test with a *p*-value < 0.05 considered statistically significant.

### 2.11 Drug target investigation

We investigated if any of the gene expression markers were previously annotated as drug targets by querying the Drug-Gene Interaction Database (DGIdb) [69] for the predicted markers using the R package rDGIdb [70,71] and only cancer-specific data sources: DoCM, JAX-CKB, MyCancerGenome, ClearityFoundationBiomarkers, MyCancerGenomeClinicalTrial, COSMIC, NCI, OncoKB, CGI, TALC, CIViC, CancerCommons, and ClearityFoundationClinicalTrial.

## 3 Results

### 3.1 Differentially expressed genes (DEGs) between B- and T-ALL subtypes

At first, we aimed to identify which ENSEMBL gene IDs are differentially expressed when comparing the B- and T-ALL subtypes in the TARGET-ALL cohort. We used three methods and four designs for DEA (see 2 Methods). Comparing intersections of identified DEGs between the three methods and four designs revealed the fewest up- and down-regulated DEGs using the year of diagnosis design in all three methods, except for the upregulated DEGs predicted by DESeq2 where the tissue portion and year of diagnosis design identified the fewest DEGs (**Table 2**). Additionally, through these comparisons, we found that the year of diagnosis design was the only one where upregulated DEGs predicted by one tool were not predicted as downregulated by another tool and vice versa (**Figure 2A**). Thus, we decided to retain the DEA performed using the year of diagnosis as a batch factor for downstream analyses. We retained only those ENSEMBL gene IDs that were in agreement as up- or downregulated according to the three methods, resulting in a set of 3848 consensus DEGs with 1729 and 2119 up- and downregulated DEGs, respectively **(Figure 2A**). We found that the log2FC values of these consensus DEGs predicted by the three DEA methods are similarly distributed **(Figure 2B)**. We did not observe any statistically significant difference in means between the log2FC values of the consensus DEGs predicted by the three DEA tools (*p*-value from oneway ANOVA = 0.251) **(Figure 2B)**. The log2FC values are interpreted as the DEGs being up- or downregulated in B-ALL compared to T-ALL.

**Figure 2.**
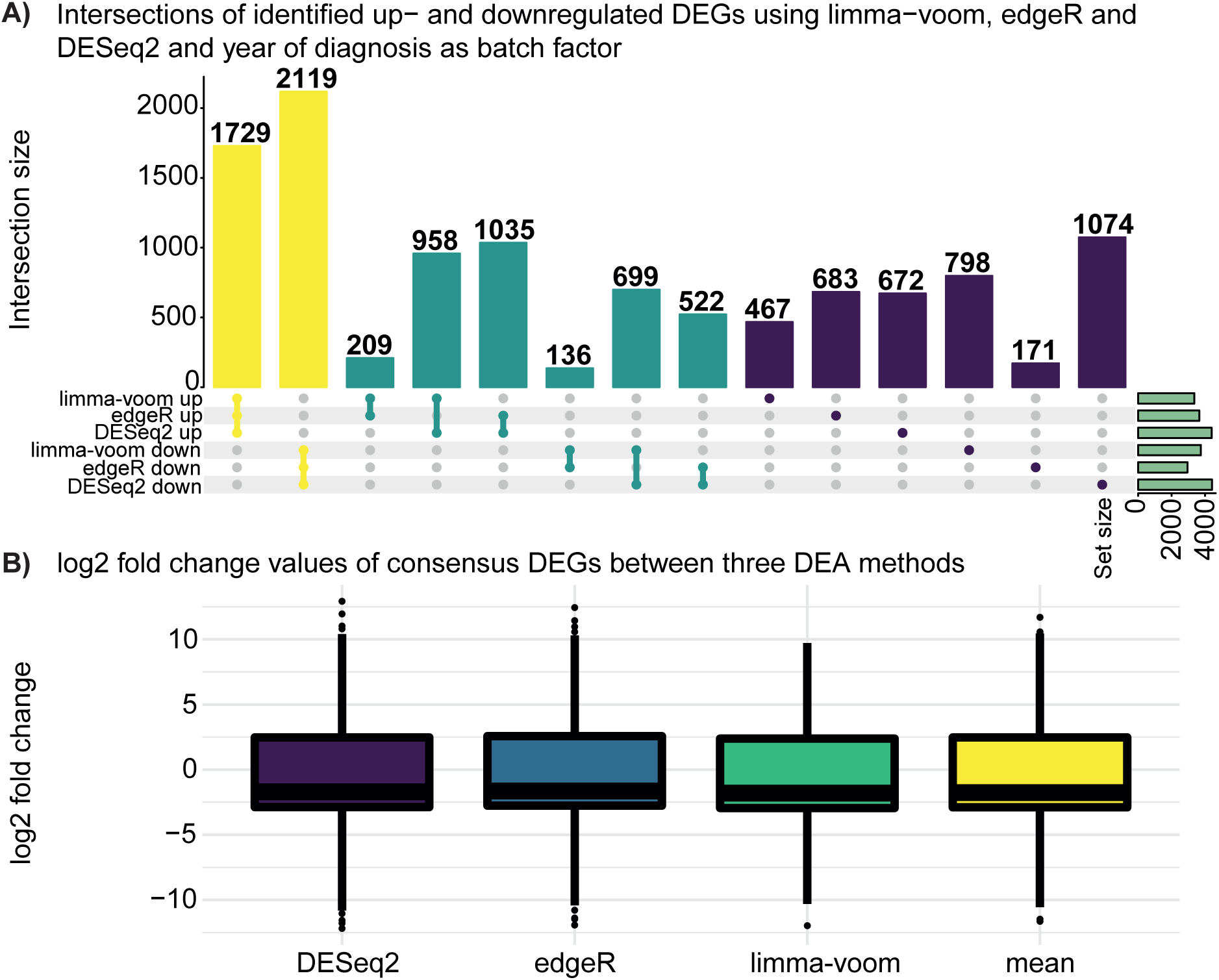
Identified up- and downregulated differentially expressed genes (DEGs) using three differential expression analysis (DEA) methods: limma-voom, edgeR, and DESeq2 with years of diagnosis as batch factor. **A)** The colors of the bars represent sets containing three (yellow), two (turquoise), or one (purple) of the up- and downregulated DEG sets identified using the three different DEA methods. The yellow and turquoise colors represent distinct overlaps between sets. Numbers above bars represent the number of identified DEGs in each intersection. Green horizontal bars to the right indicate sizes of the sets containing the up- and downregulated DEGs. See GitHub repository for similar UpSet plots with other batch factor designs in the DEA. **B)** Distribution of log2 fold change (log2FC) values of those DEGs in common between the three DEA methods: limma-voom, edgeR, and DESeq2 performed using years of diagnosis as batch factor. The distribution of the mean log2FC of these common DEGs of all three DEA methods is also shown. No statistical significant difference in means between the log2FC values of the common DEGs predicted by the three DEA tools (*p*-value = 0.251) was observed using a one-way ANOVA.

**Table 2.** Number of up- and downregulated differentially expressed genes (DEGs) identified using three methods: limma-voom, edgeR, and DESeq2 and four different designs: no batch factor, tissue portion as batch factor, tissue portion and year of diagnosis as batch factors, and year of diagnosis as batch factor.

| Design | Upregulated DEGs - limma-voom | Downregulated DEGs - limma-voom | Upregulated DEGs - edgeR | Downregulated DEGs - edgeR | Upregulated DEGs - DESeq2 | Downregulated DEGs - DESeq2 |
| --- | --- | --- | --- | --- | --- | --- |
| No batch factor | 4805 | 5356 | 3989 | 3556 | 4939 | 5416 |
| Tissue portion as batch factor | 5384 | 6087 | 4084 | 5095 | 4919 | 7935 |
| Tissue portion and year of diagnosis as batch factors | 4164 | 5054 | 3972 | 5246 | 4016 | 8003 |
| Year of diagnosis as batch factor | 3363 | 3752 | 3656 | 2948 | 4394 | 4414 |

Abbreviations: DEGs, differentially expressed genes.

As a QC of our consensus set of DEGs, we explored the presence of any reported housekeeping genes, as these are not expected to be differentially expressed. Eisenberg and Levanon (2013) provided a list of 3804 human housekeeping genes (https://www.tau.ac.il/~elieis/HKG/) expressed uniformly across 16 normal human tissue types, including white blood cells [72]. Intersecting our consensus DEGs with the list by Eisenberg and Levanon (2013) revealed an overlap of 103. We investigated the distribution of the log2FC values of the 103 housekeeping DEGs predicted by the three DEA methods (**Supplementary Figure S2**, **Supplementary Table S2**). We found that most of the 103 housekeeping DEGs are upregulated in B-ALL compared to T-ALL, with fold changes between two and 16. Next, we assessed the extent to which the housekeeping genes are dysregulated compared to the full set of genes in our dataset by calculating the ratio between the number of dysregulated housekeeping genes normalized by the total number of housekeeping genes in the dataset and the number of dysregulated genes in the dataset normalized by the total number of genes. We obtained a ratio of (103 / 3576) / (3848 / 42271) = 0.32, suggesting that the observed number of dysregulated housekeeping genes is lower than expected compared to the overall gene population.

We explored the biological roles of the up-and downregulated consensus DEGs through enrichment analysis (**Figure 3**). The consensus DEGs upregulated in B-ALL compared to T-ALL have molecular functions related to immunological activities, transforming growth factor (TGF)-beta receptor binding, and transmembrane receptor protein kinase activity (**Figure 3A**). Similarly, we also find immunological processes and transmembrane receptor protein kinase signaling overrepresented among the upregulated consensus DEGs regarding GO biological processes. Moreover, DEGs that regulate epithelial-to-mesenchymal transition are upregulated in B-ALL compared to T-ALL (**Figure 3B**). Finally, we observe that the upregulated consensus DEGs participate in various hallmarks defined by the Molecular Signatures database (MSigDB). For example, these DEGs play a role in epithelial-to-mesenchymal transition and inflammatory response, complementing the enriched GO biological process terms. These upregulated consensus DEGs are also involved in signaling pathways such as NOTCH, Wnt-beta catenin, TNF-alpha via NF-kb, and IL-2/STAT5 signaling (**Figure 3C**). NOTCH and Wnt-beta catenin pathways have previously been implicated in B- and T-ALL pathogenesis [73–77]. STAT5 activation has been found to be associated with T-ALL [78,79]. On the other hand, the consensus DEGs downregulated in B-ALL compared to T-ALL are involved in biological processes related to genome organization (**Figure 3D**). Indeed, alterations in genome organization can lead to cancer [80], and chromosomal alterations are often observed in both B- and T-ALL [81–84].

**Figure 3.**
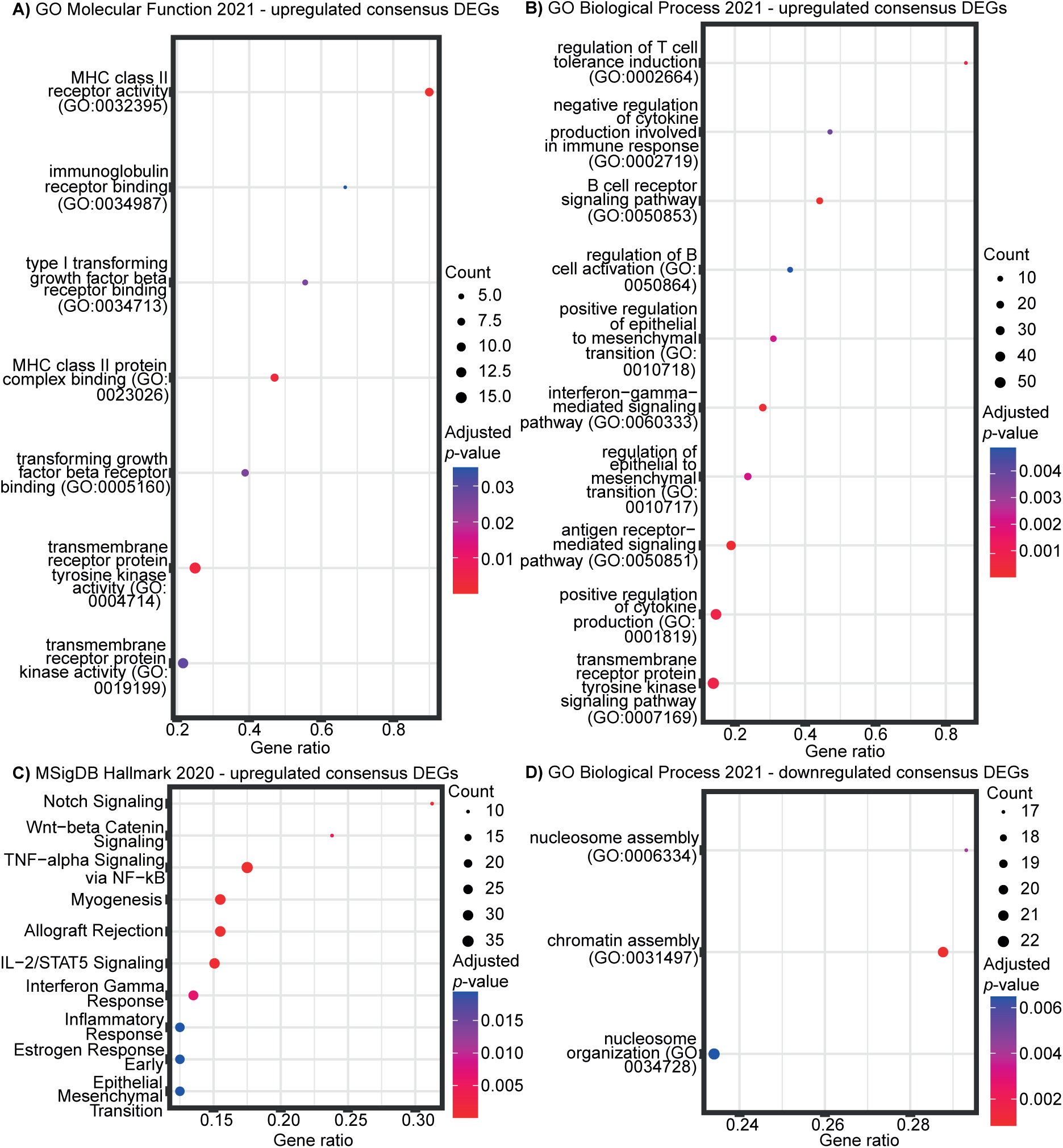
Enrichment analyses of 3848 consensus DEGs. Consensus DEGs were identified as those genes that were in agreement as up- or downregulated according to the three DEA methods: limmavoom, edgeR, and DESeq2. Enrichment analyses were performed on the 1729 upregulated consensus DEGs using the **A)** GO Molecular Function 2021 database, **B)** GO Biological Process 2021 database, and **C)** MSigDB Hallmark 2020 database, and on the 2119 downregulated consensus DEGs using the GO Biological Process 2021 database. Enrichment analyses on downregulated consensus DEGs using the GO Molecular Function 2021 and MSigDB Hallmark 2021 databases did not reveal any significantly enriched terms. In all plots, top 10 significantly enriched terms are shown (adjusted *p*-value < 0.05). Gene ratios refer to the ratio between the number of up/downregulated consensus DEGs overlapping with genes annotated in the respective term and the total number of genes annotated in the respective term. The points are colored according to adjusted *p*-value and sized according to the number of genes.

### 3.2 Definition of a minimal subset of subtype-related markers

Even upon a consensus among different methods, the DEA returned a relatively large number of DEGs (3848 DEGs). Thus, we applied two additional approaches, elastic net logistic regression and dimensionality reduction, to pinpoint candidate markers that drive the differences between B- and T-ALL. In previous work, we applied a similar approach to breast cancer subtypes, allowing us to prioritize the most important markers [59]. From elastic net logistic regression, performed on batch corrected data of the whole dataset (42271 ENSEMBL gene IDs and 378 samples), we found 31 ENSEMBL gene IDs that were selected as features in all 10 runs, comprising an intersected set of ENSEMBL gene IDs (**Figure 4A**, **Supplementary Table S3**). None of these 31 ENSEMBL gene IDs overlapped with the 103 housekeeping DEGs. We found low mean cross-validation errors in all 10 seed runs (**Figure 4B**), indicating that the trained models perform well. Elastic net regression yielded an average prediction error of 0% (no errors) across the 10 runs when predicting the 96-sample test dataset. Since the samples belonging to the two subtypes are well-separated (**Figure 4C**), we were able to train a good predictor that can classify the test data perfectly. Moreover, we are here using the model for gene selection rather than for prediction of classes. The 31 intersected ENSEMBL gene IDs predicted by elastic net logistic regression were all found to be part of the 3848 consensus DEGs. Comparing the average elastic net coefficient (average coefficient across the 10 seed runs) with the average log2FC value (average of log2FC across the three DEA methods), we found a significant negative correlation between these two values for each of the 31 ENSEMBL gene IDs (Pearson correlation coefficient: -0.8177, *p*-value: 1.9546e-8). For instance, the *BLNK* gene has an average log2FC of 7.0636, meaning *BLNK* is ∼133 times more expressed in B-ALL than in T-ALL. Further, *BLNK* has an average elastic net coefficient value of -0.0063, meaning as the expression of *BLNK* increases, the predicted class probability moves towards class 0, representing B-ALL. Finally, we examined the biotypes of the 31 ENSEMBL gene IDs and found that the largest biotype category was protein-coding genes.

**Figure 4.**
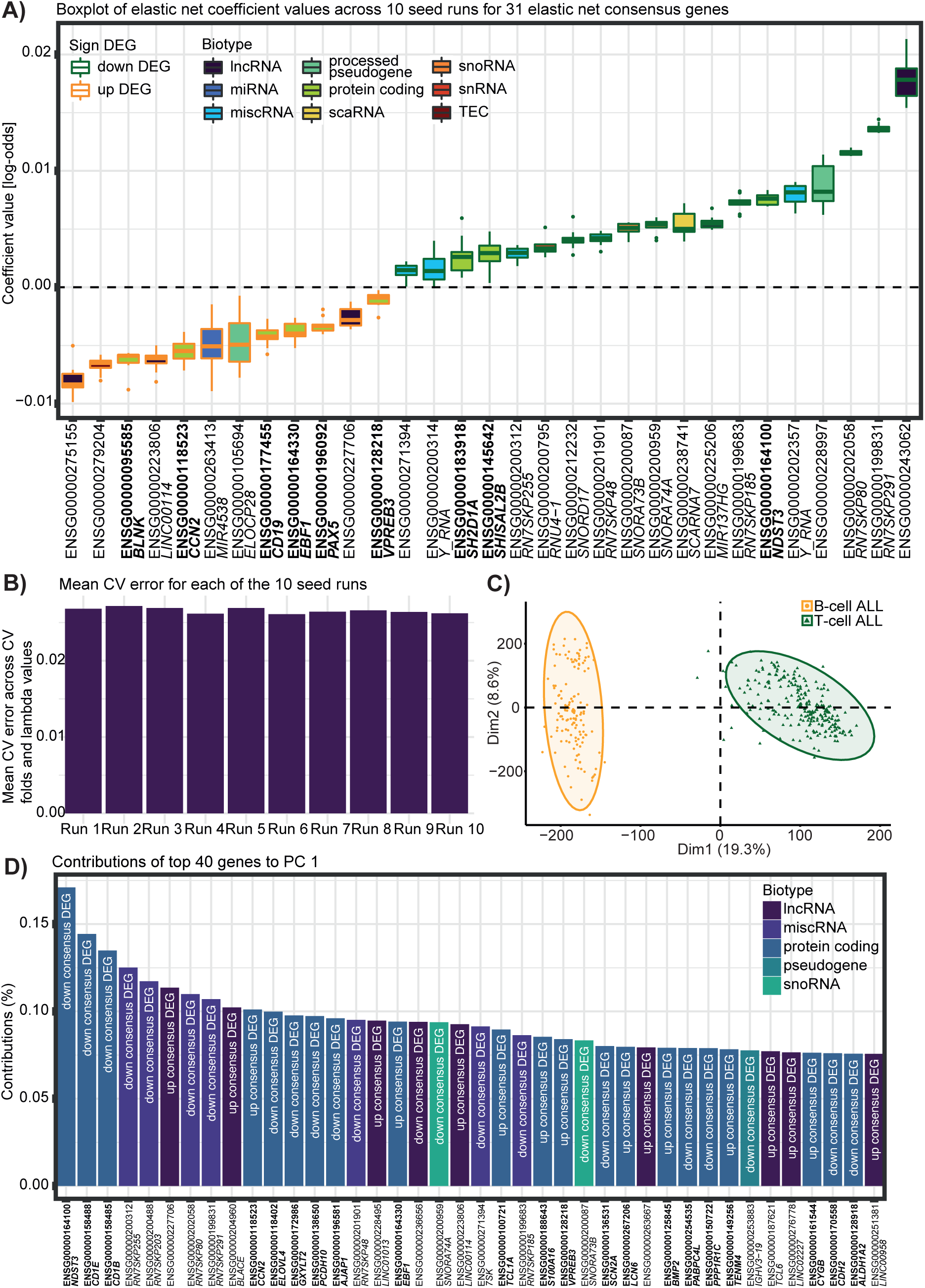
Results of elastic net logistic regression and principal component analysis (PCA). **A)** Coefficients represented as log-odds of 31 ENSEMBL gene IDs selected as features in 10 elastic net binomial logistic regression runs. Elastic net binomial logistic regression was performed on TARGET ALL batch corrected data with a dichotomized target variable encoded as 0 corresponding to B-ALL and 1 corresponding to T-ALL. The 31 ENSEMBL gene IDs are colored according to their biotype as found in the ENSEMBL database and whether they are up- or downregulated. Protein coding genes are marked in bold. The dotted horizontal line shows separation of the up- and downregulated ENSEMBL gene IDs and ENSEMBL gene IDs with negative and positive coefficients. Some ENSEMBL IDs do not have a corresponding gene name. **B)** Mean cross-validation error across cross-validation folds and lambda values for each of the 10 elastic net logistic regression where 10 different seeds have been used. **C)** PCA of TARGET ALL batch corrected data where samples are colored according to subtype. **D)** Contributions in % of top 40 ENSEMBL gene IDs contributing to PC dimension one. Contributions were found through PCA on TARGET ALL batch corrected data. The 40 ENSEMBL gene IDs are colored according to their biotype as found in the ENSEMBL database. Protein coding genes are marked in bold. For each ENSEMBL gene ID, it is indicated if it is a non-, upregulated or downregulated consensus DEG. Some ENSEMBL IDs do not have a corresponding gene name.

These protein-coding genes are *NDST3*, *BLNK*, *CCN2*, *CD19*, *EBF1*, *PAX5*, *SHISAL2B*, *SH2D1A*, and *VPREB3* (**Figure 4A**, **Supplementary Table S3**).

To further complement the results from elastic net logistic regression, we performed PCA to investigate which ENSEMBL gene IDs contribute the most towards separating the samples belonging to the two ALL subtypes. We observe that the two subtypes are mainly separated along PC1 (**Figure 4C**), which explains 19.3% of the variance in the data (**Supplementary Figure S3**). For this reason, we examined the top 40 ENSEMBL gene IDs with the highest contribution of explained variance between the ALL samples along PC1 (**Figure 4D**). None of these top 40 ENSEMBL gene IDs overlapped with the 103 housekeeping DEGs. We found that these top 40 ENSEMBL gene IDs were among the 3848 consensus DEGs and had large log2FC values (**Supplementary Table S4**), indicating that PC1 captures the highly DEGs as those contributing the most towards the separation of the two ALL subtypes. Of the top 40 ENSEMBL gene IDs, 21 are protein-coding genes: *NDST3*, *CD1E*, *CD1B*, *CCN2*, *ELOVL4*, *GXYLT2*, *PCDH10*, *AJAP1*, *EBF1*, *TCL1A*, *S100A16*, *VPREB3*, *SCN2A*, *LCN6*, *BMP2*, *PABPC4L*, *PPP1R1C*, *TENM4*, *CYGB*, *CDH2*, and *ALDH1A2* (**Figure 4D**).

### 3.3 Definition of a minimal subset of subtype-related markers across methods

We compared the ENSEMBL gene IDs discovered by consensus DEA, elastic net logistic regression, and PCA in UpSet plots (**Figure 5**). We found 14 ENSEMBL gene IDs in common between all three methods, which were not part of the 103 housekeeping DEGs: *CCN2*, *VPREB3*, *NDST3*, *EBF1*, RN7SKP185, RN7SKP291, SNORA73B, RN7SKP255, SNORA74A, RN7SKP48, RN7SKP80, LINC00114, a novel gene (ENSG00000227706), and 7SK (**Figure 5A**). Examining the biotypes of these 14 ENSEMBL gene IDs reveals two long non-coding RNA (LINC00114 and ENSG00000227706), six miscellaneous RNA (RN7SKP185, RN7SKP291, RN7SKP255, RN7SKP48, RN7SKP80, and 7SK), four protein coding (*CCN2*, *VPREB3*, *NDST3*, and *EBF1*) and two small nucleolar RNA (SNORA73B and SNORA74A). These 14 ENSEMBL gene IDs provide a minimal subset of ENSEMBL gene IDs that contribute the most towards explaining the separation observed between the two ALL cancer subtypes. Five and nine of these genes are upregulated and downregulated in B-ALL compared to T-ALL (**Table 3**).

**Figure 5.**
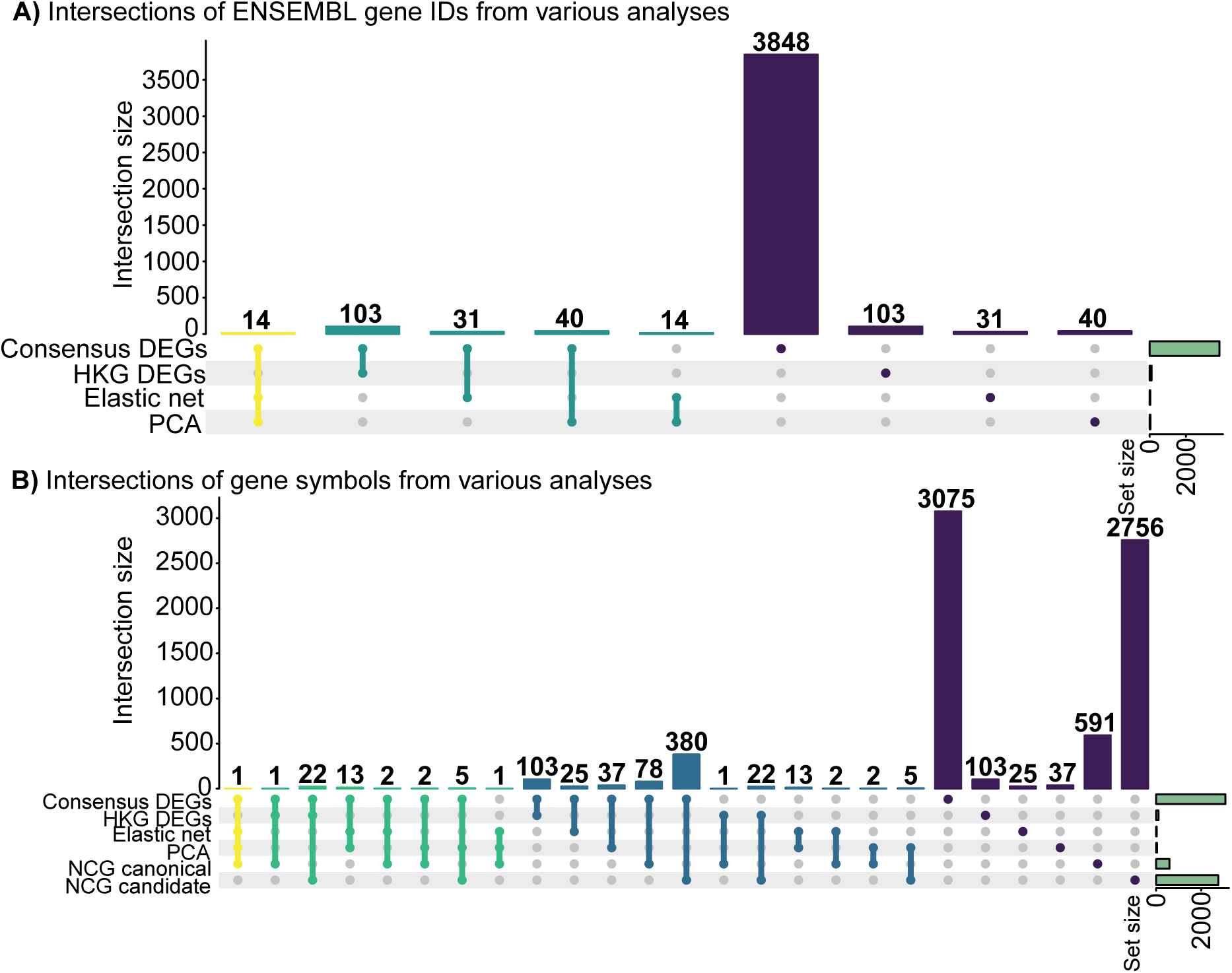
Comparison of ENSEMBL gene IDs **A)** and their external gene names **B)** discovered by consensus differential expression analysis (DEA), elastic net logistic regression, and principal component analysis (PCA). Overlap with housekeeping consensus differentially expressed genes (DEGs) **A)** and cancer genes from the Network of Cancer Genes database (NCG) **B)** are also included. In **A)**, the colors of the bars represent sets containing three (yellow), two (turquoise), or one (purple) of the ENSEMBL gene IDs identified from the different analyses. The yellow and turquoise colors represent intersections between sets. In **B)**, the colors of the bars represent sets containing four (yellow), three (green), two (blue), or one (purple) of the external gene names. The yellow, green, and blue colors represent intersections between sets. Numbers above bars represent the number of identified ENSEMBL gene IDs/external gene names in each intersection. Green horizontal bars to the right indicate sizes of the sets containing the discovered ENSEMBL gene IDs/external gene names.

**Table 3.** Average log2 fold change (log2FC) and false discovery rate (FDR) values of the defined subset of 14 subtype-related gene expression markers. The average log2FC and FDR values are across three differential expression analysis (DEA) methods (limma-voom, edgeR, and DESeq2) with the standard deviations (SD) included in brackets.

| Gene | Average log2FC [SD] | Average FDR [SD] |
| --- | --- | --- |
| LINC00114 | 8.7047 [0.2038] | 2.9924e-87 [5.1830e-87] |
| Novel gene<br>(ENSG00000227706) | 11.6906 [1.7298] | 1.0997e-90 [1.9047e-90] |
| RN7SKP185 | -7.7758 [0.6542] | 1.1201e-70 [1.5208e-70] |
| RN7SKP291 | -9.3511 [0.2172] | 5.5305e-87 [9.5791e-87] |
| RN7SKP255 | -8.1543 [1.5475] | 8.8446e-38 [1.5319e-37] |
| RN7SKP48 | -7.6915 [1.0032] | 2.3876e-47 [4.1355e-47] |
| RN7SKP80 | -8.7568 [0.7806] | 3.3216e-80 [5.7532e-80] |
| 7SK | -7.4783 [0.7991] | 3.7501e-47 [6.4834e-47] |
| CCN2 | 9.3982 [0.2464] | 2.3999e-81 [4.1568e-81] |
| VPREB3 | 9.2479 [0.4986] | 1.0546e-106 [1.8266e-106] |
| NDST3 | -11.6426 [0.3028] | 2.1470e-77 [3.7186e-77] |
| EBF1 | 8.1850 [0.7341] | 9.4078e-103 [1.6295e-102] |
| SNORA73B | -7.3992 [0.9972] | 2.1027e-44 [3.6420e-44] |
| SNORA74A | -8.6670 [0.3967] | 1.3186e-58 [2.2837e-58] |

Abbreviations: log2FC, log2 fold change; FDR, false discovery rate; DEA, differential expression analysis; SD, standard deviation.

Moreover, we compared the results of the consensus DEA, elastic net logistic regression, and PCA with the Network of Cancer Genes (NCG) database [85,86] to investigate if our results contained any genes annotated to play a role in cancer (**Figure 5B**). NCG contains two categories of cancer genes: canonical genes and candidate genes. The canonical genes have been proven experimentally to play a role in cancer. In contrast, the candidate genes contain somatic alterations predicted to play a role in cancer but lack experimental verifications [85,86]. Interestingly, we found one gene (*EBF1*) discovered in consensus DEA, elastic net logistic regression, and PCA, also annotated as a canonical cancer gene in NCG. Additionally, *PAX5* had features in common with the consensus DEA, elastic net logistic regression, and canonical cancer genes in NCG. *TCL1A* was common between the consensus DEA, PCA, and the NCG canonical genes. We also found five genes discovered by the consensus DEA and PCA and annotated as candidate cancer genes in NCG: *AJAP1*, *CD1B*, *CDH2*, *PABPC4L*, and *PCDH10*. Moreover, 78 and 380 consensus DEGs were annotated as canonical and candidate cancer genes in NCG, respectively.

### 3.4 Literature characterization of a defined subset of subtype-related gene expression markers

#### 3.4.1 Long non-coding RNAs

One study found that LINC00114 was significantly overexpressed in B-ALL patients compared to both healthy and T-ALL samples [87]. We also found that LINC00114 was significantly upregulated in B-ALL compared to T-ALL (**Table 3**). Additionally, LINC00114 has been shown to play a role in the development of colorectal cancer [88] and esophageal cancer [89]. ENSG00000227706 has been demonstrated to be associated with multiple myeloma [90] and acute myeloid leukemia [91] and overexpressed in leukemia [92,93].

#### 3.4.2 Miscellaneous RNA

RN7SKP255 was upregulated in lung adenocarcinoma compared with adjacent non-tumorous tissue [94]. RN7SKP80 has been found to play a contributing factor in distinguishing pancreatic cancer from normal tissue [95]. Overexpression of 7SK has been reported to induce apoptosis by inhibiting cell proliferation in kidney cancer [96]. 7SK was also found to be downregulated in chronic myeloid leukemia, breast, and colon cancer [97]. To our knowledge, the role of RN7SKP185, RN7SKP291, and RN7SKP48 in cancer has not been reported.

#### 3.4.3 Protein-coding genes

*CCN2* plays a role in cell proliferation, development, extracellular matrix production, migration, and adhesion [98]. This gene has previously been upregulated in B-ALL compared to control cell populations, and exclusive expression in B-ALL and not T-ALL has been reported [99]. Similarly, we found *CCN2* upregulated in B-ALL compared to T-ALL (**Table 3**). It is worth highlighting that *VPREB3* is a B-cell receptor component [100], which explains its upregulation in B-ALL compared to T-ALL (**Table 3**). Increased gene expression of this gene can activate the pro-survival phosphatidylinositol-3-OH kinase pathway [100]. Recently, another study also analyzed molecular differences between B-ALL and T-ALL and found *VPREB3* as a methylation and expression signature gene [101]. *EBF1* is a transcription factor involved in B-cell lineage specification and commitment [102], which explains its increased expression in B-ALL compared to T-ALL (**Table 3**). Deletions of *EBF1* have been found to be associated with B-ALL [102,103]. *NDST3* encodes an enzyme that plays a role in heparan sulfate metabolism [104]. Heparan sulfate is a glycosaminoglycan expressed on cell surfaces and in the extracellular matrix [105], which on tumor cell surfaces can promote tumorigenesis by regulating autocrine signaling resulting in uncontrolled cell growth [106]. Recently, Hu et al. (2022) found *NDST3* to correlate significantly with overall survival in acute myelogenous leukemia [107].

#### 3.4.4 Small nucleolar RNA

High expression of SNORA74A has been associated with a shorter progression free survival in chronic lymphocytic leukemia [108]. Moreover, SNORA74A has been reported as a potential oncogene in gastric cancer [109] and as a novel noninvasive diagnostic biomarker in pancreatic cancer [110]. SNORA73B was used for creating a prognostic signature together with 13 other snoRNAs, which could divide patients with acute myeloid leukemia into high- and low-risk groups [111]. In other cancer types, SNORA73B has been shown to promote development of endometrial cancer as a potential oncogene with increased expression [112], and Liu et al. (2020) created a prognostic signature based on expression values of four snoRNAs including SNORA73B in patients with sarcoma [113].

### 3.5 Prognostic potential of subtype-related gene expression markers

T-ALL carries a less favourable outcome compared to B-ALL with a 5-10% lower outcome. Reasons for this difference include older age, lower chemotherapy tolerance, less favourable low-risk genetic subtypes, higher resistance to chemotherapeutic drugs, and lower availability of targeted therapies of T-ALL compared to B-ALL [114]. To evaluate the prognostic potential of the defined subset of 14 subtype-related gene expression markers, we performed survival analyses. First, we conducted survival analysis using a multivariate Cox regression model where we included the age and sex of patients as covariates. From these analyses, we found that the expression level of all 14 markers significantly affected survival (**Table 4**). Investigating the ranking of the hazard ratios revealed that the four protein-coding genes (*VPREB3*, *EBF1*, *CCN2*, and *NDST3*) and the two long non-coding RNA (LINC00114 and ENSG00000227706) had the highest hazard ratios. In contrast, the miscellaneous RNA and the small nucleolar RNA had the lowest hazard ratios. Moreover, *VPREB3*, *EBF1*, *CCN2*, LINC00114, and ENSG00000227706 all had hazard ratios above 1 ranging between 1.22 and 1.33, indicating that a one-unit increase in expression of each of these markers is associated with a 22-33% increase in the hazard of experiencing death. On the other hand, the remaining markers had hazard ratios below 1, indicating that a one-unit increase in gene expression is associated with a decrease in the hazard of experiencing death. These results suggest a prognostic potential of the 14 gene expression markers and a greater prognostic impact of the protein-coding genes and the long non-coding RNAs compared to the miscellaneous RNA and the small nucleolar RNA.

**Table 4.** Hazard ratios of the defined subset of 14 subtype-related gene expression markers together with 95% confidence intervals and *p*-values. Hazard ratios were found from a multivariate Cox regression model with gene expression as the explanatory variable and survival data as the response variable. The model included age and sex of patients as covariates. The table is sorted by descending hazard ratios of the expression variable.

| Gene | Hazard ratio [95% CI] | *p*-value |
| --- | --- | --- |
| ***VPREB3*** | 1.33 [1.24-1.42] | 2.27e-17 |
| ***EBF1*** | 1.28 [1.20-1.37] | 4.16e-14 |
| **LINC00114** | 1.24 [1.17-1.31] | 5.92e-14 |
| **Novel gene**<br>**(ENSG00000227706)** | 1.23 [1.17-1.29] | 1.43e-17 |
| ***CCN2*** | 1.22 [1.16-1.29] | 4.68e-13 |
| ***NDST3*** | 0.848 [0.811-0.886] | 2.30e-13 |
| **SNORA74A** | 0.802 [0.757-0.850] | 9.32e-14 |
| **RN7SKP255** | 0.787 [0.746-0.831] | 2.63e-18 |
| **RN7SKP291** | 0.776 [0.728-0.827] | 6.02e-15 |
| **RN7SKP80** | 0.776 [0.730-0.824] | 2.29e-16 |
| **7SK** | 0.770 [0.725-0.817] | 1.05e-17 |
| **RN7SKP185** | 0.764 [0.718-0.813] | 3.19e-17 |
| **RN7SKP48** | 0.751 [0.703-0.801] | 6.75e-18 |
| **SNORA73B** | 0.750 [0.702-0.801] | 8.00e-18 |

Abbreviations: CI, confidence interval.

Afterwards, we also performed a Kaplan-Meier survival analysis to compare differences in survival between patients with high and low expression of each marker. We found that all 14 markers had a significant difference in survival when comparing these two groups. The Kaplan-Meier survival plots show that having high expression of *VPREB3*, *EBF1*, *CCN2*, LINC00114, and ENSG00000227706 results in lower survival probability and thus, a worse prognosis (**Figure 6**). Furthermore, these five markers were upregulated in B-ALL patients compared to T-ALL patients (**Table 3**), suggesting a worse prognosis for patients with B-ALL. In contrast, patients with a low expression of the remaining nine markers have a lower survival probability than patients with high expression (**Supplementary Figure S4**). These nine genes were downregulated in B-ALL patients compared to T-ALL patients (**Table 3**), again indicating a worse prognosis for patients with B-ALL.

**Figure 6.**
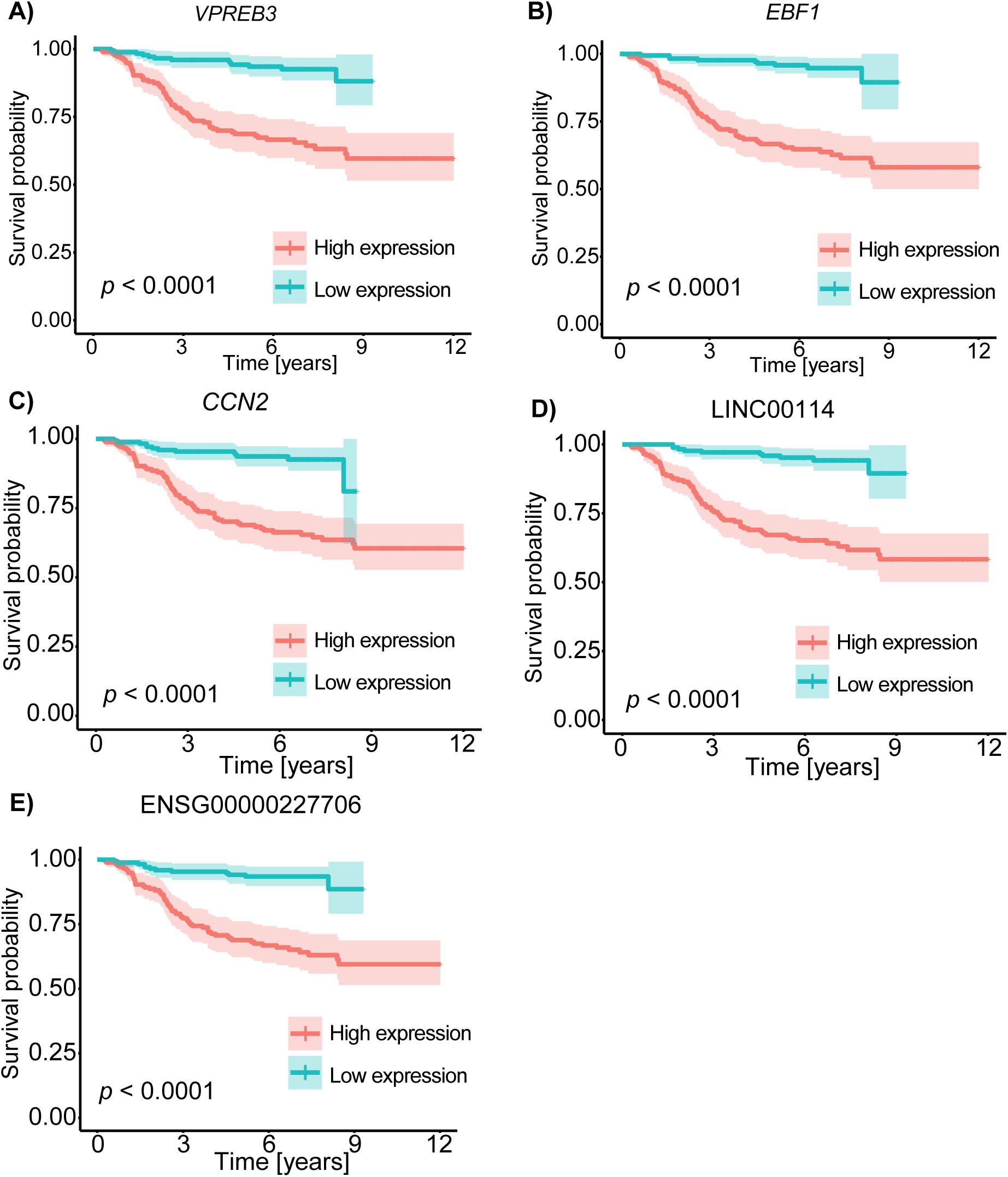
Kaplan-Meier survival plots of five of the discovered subtype-related gene expression markers. The five markers shown are three protein-coding genes: **A)** *VPREB3*, **B)** *EBF1*, **C)** *CCN2* and two long non-coding RNA: **D)** LINC00114 and **E)** ENSG00000227706. Patients were categorized into high (orange) and low (blue) expression groups based on whether their expression values were above or below the median expression level of the corresponding gene. Survival curves were constructed using the discrete expression group as the independent variable, and the significance of the difference in survival between the two groups was assessed using a log-rank test with a *p*-value < 0.05 considered statistically significant.

### 3.6 Drug target investigation

We investigated the therapeutic potential of the 14 subtype-related gene expression markers by querying these genes in the Drug Gene Interaction Database (DGIdb). One of these genes, *CCN2*, was previously annotated to interact with 17 drugs: 2−methoxyestradiol, acridine, androstanolone, curcumin, digoxin, enalapril, estradiol, inositol, insulin, liothyronine sodium, prasterone, propranolol, ramipril, spironolactone, staurosporine, thrombin, vitamin E. However, we did not find convincing literature about these drug interactions with *CCN2* in cancer.

### 3.7 Stratification of the ALL samples beyond the B- and T-ALL subtypes

To further explore the existence of subtypes within the two main ALL subtypes, we performed unsupervised clustering of the gene expression data using the *cola* framework [68]. For the optimal selection of the subgrouping, we examined 20 different clustering methods consisting of combinations of four feature selection and five partitioning methods with *k* number of clusters ranging from 2 to 6. We first compared the performance of two data inputs representing two stages of data processing: 1) batch-corrected data and 2) raw data where replicates have been adjusted for and subsequently adjusted using *cola’s* processing. We compared the predicted clusters for these two data inputs using *k* = 2 for all 20 methods with the actual subtype labels of B- and T-ALL (**Figure 7A-B**). For the batch-corrected data, 10 of the 20 methods could not 100% correctly cluster the B- and T-ALL samples into their clusters. For example, the method ATC:hclust clusters the 133 B-ALL samples into two different clusters divided into 107 samples in one cluster and 26 samples in the second cluster (**Figure 7A**). On the other hand, none of the methods could 100% correctly cluster the B- and T-ALL samples into two separate clusters when using the raw *cola*-adjusted data as input (**Figure 7B**) and in general resulted in worse classification performance. Thus, we proceeded with the batch-corrected data for further analyses.

**Figure 7.**
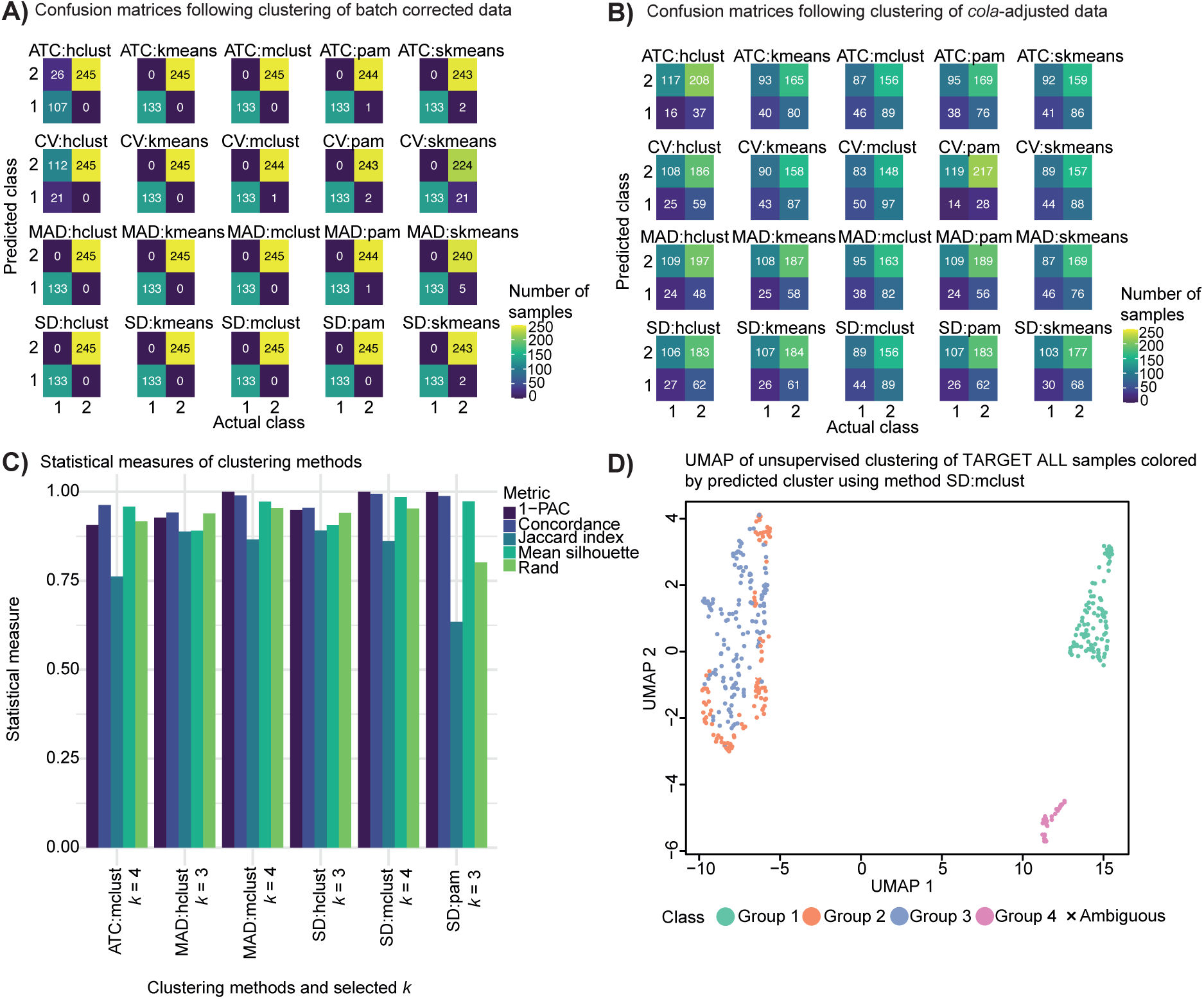
Results of unsupervised clustering. **A-B)** Confusion matrices of unsupervised clustering using the *cola* framework with *k* = 2 performed on TARGET ALL gene expression data. The unsupervised clustering was conducted using 20 different methods and subsequently compared with actual class labels. The actual class labels refer to the two annotated ALL subtypes where B- and T-ALL are encoded as 1 and 2, respectively. Each confusion matrix contains the result from each method. Values in the confusion matrices represent the number of samples. The input data was **A)** TARGET ALL batch corrected data and **B)** TARGET ALL raw data adjusted using *cola*’s processing method. **C)** Statistical measures of six unsupervised clustering methods using the *cola* framework. The six methods were chosen as those that could 100% correctly cluster the B- and T-ALL samples into two separate clusters and which do not suggest *k* = 2 as the best *k*. The unsupervised clustering was performed on TARGET ALL batch corrected data. For each method, the suggested *k* is shown. The statistical measures are 1-PAC, concordance, Jaccard index, mean silhouette, and rand. **D)** UMAP visualization of predicted clusters using method, SD:mclust. The unsupervised clustering was performed using the *cola* framework. The colored dots represent predicted class labels and the black cross represents samples with a silhouette score < 0.5.

Next, we examined the 10 methods that could 100% correctly cluster the B- and T-ALL samples into two separate clusters to find the optimal method and *k*. These methods are ATC:kmeans, ATC:mclust, CV:kmeans, MAD:hclust, MAD:kmeans, MAD:mclust, SD:hclust, SD:kmeans, SD:mclust, and SD:pam. Inspecting the suggested best *k* for these 10 methods, we observe that four methods suggest *k* = 2. Since we seek to stratify the samples beyond two subtypes, we discard these four methods suggesting *k* = 2. Finally, following inspection of the reported statistical measures of the remaining six methods (**Figure 7C**), we select the top method with the highest statistical measures, SD:mclust, which suggests *k* = 4. This method shows highly stable subgrouping for *k* = 4 (**Supplementary Figure S5**).

Visualizing the clustering using UMAP revealed a separation between three clusters: one cluster consisting of samples predicted as belonging to group 1, one cluster consisting of samples predicted as belonging to group 4, and one cluster consisting of samples predicted as belonging to group 2 and 3 (**Figure 7D**). Clusters 1 and 4 are original B-ALL subtype samples. In contrast, clusters 2 and 3 are original T-ALL subtype samples, showing that the clustering split each subtype into two further groups. To better understand the differences within each of the two subtypes (B- and T-ALL), we performed feature selection using random forest on full gene expression data and cluster label as the target classification variable. We built a random forest model separately for clusters 1 and 4 (B-ALL) and clusters 2 and 3 (T-ALL). Following 10 random forest seed runs on the predicted clusters 1 and 4, we did not find any overlap of selected ENSEMBL gene IDs (**Supplementary Table S5**). On the other hand, eight genes were selected in all 10 seed runs when applying random forest on the predicted clusters 2 and 3: *PLXND1*, *TFAP2C*, *BEX2*, *PCDH19*, *C14orf39*, *SIX6*, *MAML3*, and *SALL4P7*. The first seven genes are protein-coding genes, whereas *SALL4P7* is a transcribed processed pseudogene. None of these eight genes showed significant results from multivariate Cox regression or Kaplan-Meier survival analyses. Moreover, these eight genes have not previously been annotated as drug targets in DGIdb.

To highlight a few of these genes, *PLXND1* and *BEX2* have previously been reported as DEGs between CpG Island Methylator Phenotype (CIMP) subgroups of pediatric patients with T-ALL [115]. Furthermore, *PLXND1* has been found to be a transcriptional target of the NOTCH signaling pathway [116], and *BEX2* has been suggested as a tumor suppressor gene in glioma [117]. Similarly, one study found an association between T-ALL oncogenic subgroups and ectopic expression of a set of genes, including *SIX6* and *TFAP2C*, suggesting that abnormal expression of these genes is involved in T-ALL oncogenesis [118].

### 3.8 In silico validation of predicted gene expression markers in independent Danish cohort

To evaluate the robustness of our results, we validated the predicted gene expression markers in an independent Danish cohort of pediatric patients with ALL. This cohort consisted of 88 patients divided into 77 and 11 B- and T-ALL samples, respectively. For this analysis, we first performed a PCA to investigate if the candidate gene expression markers could separate the two ALL subtypes. We observed that the 14 markers demonstrated a more effective separation between the two subtypes (**Figure 8A**) compared to the differentiation achieved by utilizing all genes in the expression data (38710 genes) (**Figure 8B**).

**Figure 8.**
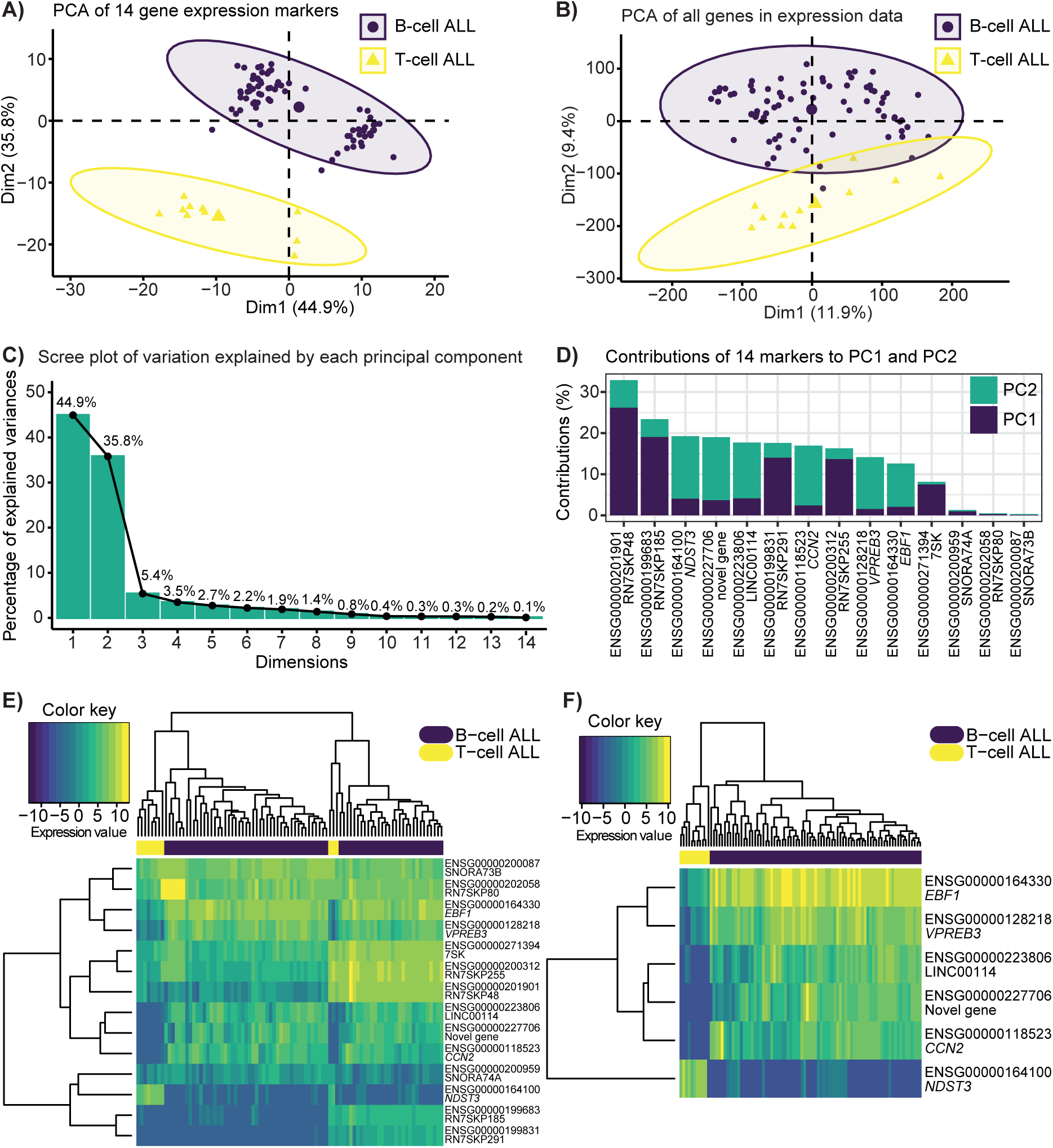
In silico validation of predicted gene expression markers using gene expression data of an independent Danish cohort of pediatric patients with ALL. **A)** PCA of expression data of the 14 markers of the Danish cohort. These 14 markers were found to distinguish the two ALL subtypes, B- and T-cell ALL, in the TARGET ALL discovery dataset. **B)** PCA of expression data of all genes in the expression dataset of the Danish cohort. **C)** Scree plot of percentage of explained variance for the 14 principal component (PC) dimensions from PCA performed in A). The percentage of explained variance for each of the 14 dimensions are shown on top of each bar. **D)** Contributions in % of the 14 markers to PC1 and PC2. **E)** Unsupervised hierarchical clustering of expression data of the 14 markers visualized as a heatmap. **F)** Unsupervised hierarchical clustering of expression data of a subset of the 14 markers visualized as a heatmap. In **E-F)**, the samples are annotated with subtype (B- and T-cell ALL) labels and values in the heatmaps are voom transformed processed expression data.

The first two PCs explain most of the variation in the data (**Figure 8C**), and investigating the contributions of each predicted gene expression marker to these two PCs demonstrate that roughly half of these (RN7SKP48, RN7SKP185, RN7SKP291, RN7SKP255, and 7SK) contribute the most to the variation observed along PC1 whereas *NDST3*, ENSG00000227706 (novel gene), LINC00114, *CCN2*, *VPREB3*, and *EBF1* contribute the most to the variation observed along PC2 (**Figure 8D**). We also investigated the top 50 genes contributing to PC1 and the top 50 genes contributing to PC2 from the PCA performed on all genes in the expression data of the Danish cohort. This revealed that three of the 14 candidate expression markers were part of the top 50 genes contributing to PC1 (RN7SKP48, RN7SKP185, and RN7SKP291). Another five of the 14 candidate expression markers belonged to the top 50 genes contributing to PC2 (LINC00114, *NDST3*, ENSG00000227706 (novel gene), *CCN2*, and *VPREB3*) (**Supplementary Figure S6**). Next, we performed unsupervised hierarchical clustering of the expression data of the predicted 14 markers (**Figure 8E**). These results revealed that *EBF1*, *VPREB3*, LINC00114, ENSG00000227706 (novel gene), *CCN2*, and *NDST3* seem to be able to separate the two ALL subtypes based on expression levels. Interestingly, these six markers were also the ones with the highest hazard ratios of the survival analyses performed on the TARGET discovery dataset (**Table 4**). Moreover, these six markers were also the ones showing the highest contribution to the observed variance along PC2 (**Figure 8D**). Indeed, the PCA (**Figure 8A**) illustrates that the two ALL subtypes are mainly separated along PC2. In accordance with this, unsupervised hierarchical clustering of expression data of these six markers demonstrates their ability to perfectly separate the two ALL subtypes (**Figure 8F**). This is in contrast to using all 14 markers where three T-ALL samples are clustered more similar to B-ALL samples than the remaining T-ALL samples (**Figure 8E**). We also validated the prognostic effect of the 14 markers in this independent cohort using Cox proportional hazards regression. Of those markers complying with the proportional hazards assumption (12 out of 14), the expression of these markers did not show significant effects on survival of the patients at the univariate level (**Supplementary Table S6**). This can be due to the fact that only five of the patients in our validation cohort have deceased, making it difficult to assess the effect.

## 4 Discussion

In this study, we have analyzed gene expression data for the prediction of gene expression markers separating two ALL subtypes, B- and T-ALL. Identifying markers differentiating ALL subtypes is important for diagnostic and prognostic purposes. For instance, one study found that expression of a circulating microRNA may be used as a non-invasive biomarker for diagnosing and predicting prognosis in pediatric patients with ALL [119]. Similarly, Wang and Zhang (2020) found that low expression of LEF1 is a biomarker of an aggressive subtype of T-ALL called early T-cell precursor, suggesting that including LEF1 with traditional immunephenotyping can enhance diagnosis of early T-cell precursor [120]. In B-ALL, one study demonstrated high and subtype-specific expression of IGF2BP3 associated with good outcome in high-risk patients, suggesting that IGF2BP3 could improve stratification and prognosis of B-ALL [121]. Finally, Cavalcante and coworkers (2016) found a set of glycoproteins as candidate biomarkers for early diagnosis of B-ALL and which may be useful to determine response to treatment [122].

In order to identify gene expression markers that can differentiate B-ALL and T-ALL, we analyzed gene expression data of an ALL cohort from TARGET, applying various approaches such as DEA and machine learning. Reliable results are dependent on proper processing of expression data. For this purpose, we established a bioinformatics processing workflow (**Figure 1**) which we showed to successfully distinguish the two ALL subtypes (**Figure 7A-B**). Indeed, gene expression data has previously been used for similar purposes. For example, Walter et al. (2021) found that whole transcript sequencing could be used to reliably classify ALL patients [26].

We discovered a small subset of ALL subtype-related gene expression markers comprised of *CCN2*, *VPREB3*, *NDST3*, *EBF1*, RN7SKP185, RN7SKP291, SNORA73B, RN7SKP255, SNORA74A, RN7SKP48, RN7SKP80, LINC00114, a novel gene (ENSG00000227706), and 7SK. These markers encompass various biotypes: long non-coding RNA, miscellaneous RNA, protein-coding genes, and small nucleolar RNA. We validated the classification ability of these markers in an independent cohort of Danish patients with ALL and found that a subset of these 14 markers (*EBF1*, *VPREB3*, LINC00114, ENSG00000227706, *CCN2*, and *NDST3*) could perfectly separate B- and T-ALL in this independent cohort (**Figure 8**). While extensive characterizations of these markers in ALL are less established, the majority of them (LINC00114, novel gene (ENSG00000227706), RN7SKP255, RN7SKP80, 7SK, *CCN2*, *VPREB3*, *EBF1*, *NDST3*, SNORA74A, and SNORA73B) have previously been implicated in other cancer types including other leukemia types as described in section 3.4. The four protein-coding markers (*CCN2*, *VPREB3*, *EBF1*, and *NDST3*) and 7SK have been described to play a role in various cellular pathways such as apoptosis, cell proliferation, and survival. A subset of these markers has also previously been implicated in differences between B- and T-ALL. For example, LINC00114 and *CCN2* have previously been found to be upregulated in B-ALL compared to T-ALL while deletions of *EBF1* have been associated with B-ALL [102,103] and *VPREB3* has been found as a methylation and expression signature gene between B- and T-ALL [101]. Comparing the different biotypes of the predicted markers, the protein-coding genes are described the most in literature. This is likely due to their well-established biological roles and a greater historical focus on protein-coding genes than other gene types such as non-coding RNAs. Additionally, protein-coding genes encode functional protein products that play a role in various signaling pathways, making them notable targets for further exploration. Nevertheless, in the past few decades, non-coding RNAs have received increasing recognition for their roles in cancer [123–127].

Following definition of this small subset of subtype-related expression markers, we evaluated their prognostic and therapeutic potential. We found that the expression level of all 14 markers had a prognostic effect on the survival of the patients. In particular, we found that high expression of *VPREB3*, *EBF1*, *CCN2*, LINC00114, and ENSG00000227706 (novel gene) and low expression of *NDST3*, RN7SKP185, RN7SKP291, SNORA73B, RN7SKP255, SNORA74A, RN7SKP48, RN7SKP80, and 7SK resulted in lower survival probability. The first five markers were all upregulated in B-ALL compared to T-ALL and the remaining nine markers were downregulated in B-ALL compared to T-ALL, suggesting a worse prognosis for patients with B-ALL. Additionally, we found that one of these markers, *CCN2*, had previously been reported as a drug target in DGIdb [69]. Considering the multifaceted role of *CCN2* in cancer, modulating its activity could be explored for therapeutic purposes. Given the upregulation of *CCN2* in B-ALL compared to T-ALL, targeting *CCN2* may offer a strategy to mitigate aberrant cellular processes in ALL such as cell proliferation, migration, and adhesion.

We also clustered the expression data to predict further subgroups beyond the two major ALL subtypes. We discovered four clusters that separated the B-ALL samples into two clusters and the T-ALL samples into two clusters. We found eight genes driving separation between the two predicted clusters of the T-ALL samples: *PLXND1*, *TFAP2C*, *BEX2*, *PCDH19*, *C14orf39*, *SIX6*, *MAML3*, and *SALL4P7*. The majority of these have previously been described to play a role in cancer. Various studies have grouped patients with ALL into multiple subtypes beyond B- and T-ALL, and further genetic subtypes have been proposed within B-ALL which are associated with patient prognosis. For example, Li et al. (2018) defined 14 gene expression subgroups where eight of them were also previously described. These subgroups are characterized by gene fusions, hyperdiploidy, and mutations in specific genes [22]. In contrast to B-ALL, genetic subtypes with clinical relevance have not yet been clearly established in T-ALL [7,114]. Nevertheless, studies have classified T-ALL into multiple subgroups. For example, Liu et al. (2017) identified eight subgroups of patients with T-ALL based on genetic alterations and aberrant expression of various transcription factors [23]. Stratifying patients with ALL into novel subgroups is of clinical value as this can aid disease classification, guide targeted therapies, inform prognosis, and facilitate risk stratification [20,22].

One of our applied methods for discovery of gene expression markers was elastic net logistic regression which resulted in an average prediction error of 0% when predicting the test dataset. This is attributed to the already well-separated dataset (**Figure 4C**). Here, it is worth noting that we are not relying on the results of elastic net logistic regression alone but as part of a collection of multiple analyses that together serve to pinpoint candidate markers driving the differences between the two ALL subtypes. Indeed, this study has taken an ensemble approach combining results from multiple methods to increase confidence in the predicted results. For instance, we created consensus DEA results across three DEA methods. This approach has previously been reported to generate a list of DEGs with great accuracy, indicating that combining various methods can produce more suitable results [128]. Moreover, we applied different machine learning approaches to discover subtype-related markers across these methods, and furthermore, we intersected results from elastic net logistic regression and random forest across 10 seed runs. Ensemble machine learning has previously been reported to outperform single classifiers. For example, Xiao et al. (2018) used deep neural networks to ensemble five machine learning classification models for cancer prediction which resulted in more accurate prediction than the single classifiers [129]. A limitation of this study is the lack of normal control samples, rendering comparison between the two ALL subtypes challenging as these subtypes originate from different cell types. While healthy tissue RNA-seq data is available from e.g. Genotype-Tissue Expression Portal [130], we could not find a certain source for specifically a children normal tissue dataset, which is important to not bias the analysis as adult and pediatric ALL have been shown to exhibit differences [131,132].

Future investigation is needed to elucidate the mechanisms of the deregulation of the predicted expression markers including comparisons with normal controls, coupled with mechanistic evidence such as mutations, epigenetic aberrations or chromosomal rearrangements. The future of ALL research likely continues increasing our molecular knowledge of ALL and identifying novel markers for early detection, prognosis, and treatment evaluation, with the ultimate goal of integrating these into clinical practice to enhance ALL management.

## 5 Conclusion

In this study, we discovered 14 candidate gene expression markers separating the two main ALL subtypes (B- and T-ALL), important for diagnostic and prognostic purposes. We found that the expression levels of these 14 markers had significant effects on survival of the patients, suggesting worse prognosis for B-ALL. Stratifying patients with ALL into further subgroups is crucial for improving disease classification, guiding targeted therapies, and facilitating risk stratification, ultimately enhancing clinical decision-making. Here, we discovered four clusters with eight genes driving separation between two of these clusters. Further research is needed to investigate the mechanisms of the deregulation of the predicted markers by incorporating evidence of mutations, epigenetic changes or chromosomal rearrangements.

## Abbreviations

1-PAC: 1-the proportion of ambiguous clustering
ALL: acute lymphoblastic leukemia
ATC: ability to correlate to other rows
B-ALL: B-cell precursor acute lymphoblastic leukemia
CIMP: CpG Island Methylator Phenotype
CV: coefficient of variation
DEA: differential expression analysis
DGIdb: Drug-Gene Interaction Database
DEG: differentially expressed gene
FDR: false discovery rate
hclust: hierarchical clustering
kmeans: k-means clustering
log2FC: log2 fold change
MAD: median absolute deviation
MDS: multidimensional scaling
MSigDB: Molecular Signatures database
mclust: model-based clustering
NCG: Network of Cancer Genes
OOB: out-of-bag
PCA: principal component analysis
PC: principal component
pam: partitioning around medoids
QC: quality control
RNA-seq: RNA sequencing
SD: standard deviation
skmeans: spherical k-means clustering
TARGET: Therapeutically Applicable Research to Generate Effective Treatments
TCGA: The Cancer Genome Atlas
T-ALL: T-cell acute lymphoblastic leukemia
TGF: transforming growth factor

## Acknowledgements

This project is supported by The European Union’s Interregional Öresund–Kattegat–Skagerrak grant. This work is part of Interregional Childhood Oncology Precision Medicine Exploration (iCOPE), a cross-Oresund collaboration between University Hospital Copenhagen, Rigshospitalet, Lund University, Region Skåne and Technical University Denmark (DTU), supported by the European Regional Development Fund. This project is supported by Elegant North (EN; Exploring Leukemia: Education Genetics And Technology; New Option for Rare diseases Towards Health), a collaboration between Oslo University Hospital, Capital Region of Denmark, Technical University Denmark (DTU), Abzu, Plesner, Region Skåne. This project is also supported by Danmarks Grundforskningsfond (Grant/Award Number: DNRF125). The results published here are in part based upon data generated by the Therapeutically Applicable Research to Generate Effective Treatments (https://www.cancer.gov/ccg/research/genome-sequencing/target) initiative, phs000218. The data used for this analysis are available at the Genomic Data Commons (https://portal.gdc.cancer.gov). dbGaP Sub-study ID: phs000464. The authors would like to acknowledge Marianne Helenius, Christian Højte Schouw, and Lars Rønn Olsen for their valuable insight and helpful discussions.

## Data accessibility

The data that support the findings of this study are openly available in the Therapeutically Applicable Research to Generate Effective Treatments (https://www.cancer.gov/ccg/research/genome-sequencing/target) initiative, phs000218. The data used for this analysis are available at the Genomic Data Commons (https://portal.gdc.cancer.gov). dbGaP Sub-study ID: phs000464. GitHub and OSF repositories associated with this study are available at https://github.com/ELELAB/ALL_markers, https://github.com/ELELAB/RNA_DE_pipeline, and https://osf.io/kgfpv/.

## Author contributions

MN contributed with conceptualization, investigation, methodology, code development, data interpretation, visualization, and writing of original draft and review and editing. NT contributed with code development, investigation, data interpretation, and writing of original draft and review and editing. ASL contributed with code development, investigation, and data interpretation. HBLP contributed with code development. UKS contributed with data acquisition. KW contributed with data acquisition and funding acquisition. KS contributed with data acquisition and funding acquisition. MT contributed with conceptualization, investigation, methodology, data interpretation, supervision, and writing of original draft and review and editing. EP contributed with conceptualization, data curation, funding acquisition, investigation, methodology, data interpretation, project administration, resources, supervision, and writing of original draft and review and editing.

## Supporting Information

**Supplementary_Table_S1.xlsx**

**Information of lost genes when using the updated geneInfoHT table in TCGAbiolinks containing GC content annotations.**

**Supplementary_Table_S2.pdf**

**List of 103 housekeeping differentially expressed genes.**

**Supplementary_Table_S3.pdf**

**Consensus set of selected ENSEMBL gene IDs from elastic net binomial logistic regression.**

**Supplementary_Table_S4.pdf**

**Top 40 ENSEMBL gene IDs with highest contribution of explained variance between acute lymphoblastic leukemia (ALL) samples along the first principal component.**

**Supplementary_Table_S5.pdf**

**Selected ENSEMBL gene IDs from random forest variable selection.**

**Supplementary_Table_S6.pdf**

**Hazard ratios of the defined subset of 14 subtype-related gene expression markers.**

**Supplementary_Text_S1.pdf**

**Details on updated GC content and gene length annotations in TCGAbiolinks.**

**Supplementary_Text_S2.pdf**

**Details of pipeline settings of RNA sequencing pipeline.**

**Supplementary_Figure_S1.pdf**

**Overview of the TARGET ALL cohort.**

**Supplementary_Figure_S2.pdf**

**Density plot of log2 fold change values of 103 housekeeping consensus differentially expressed genes.**

**Supplementary_Figure_S3.pdf**

**Scree plot of percentage of explained variance for the first 20 principal component dimensions from principal component analysis.**

**Supplementary_Figure_S4.pdf**

**Kaplan-Meier survival plots of nine of the discovered subtype-related gene expression markers.**

**Supplementary_Figure_S5.pdf**

**Visualization of unsupervised clustering using method SD:mclust. Supplementary_Figure_S6.pdf**

**Contributions in % of the top 50 genes to principal component 1 and principal component 2 performed on gene expression data of a Danish cohort of pediatric patients with ALL.**

