## Supplementary Figure S1 for "Data-driven discovery of gene expression markers distinguishing pediatric acute lymphoblastic leukemia subtypes"

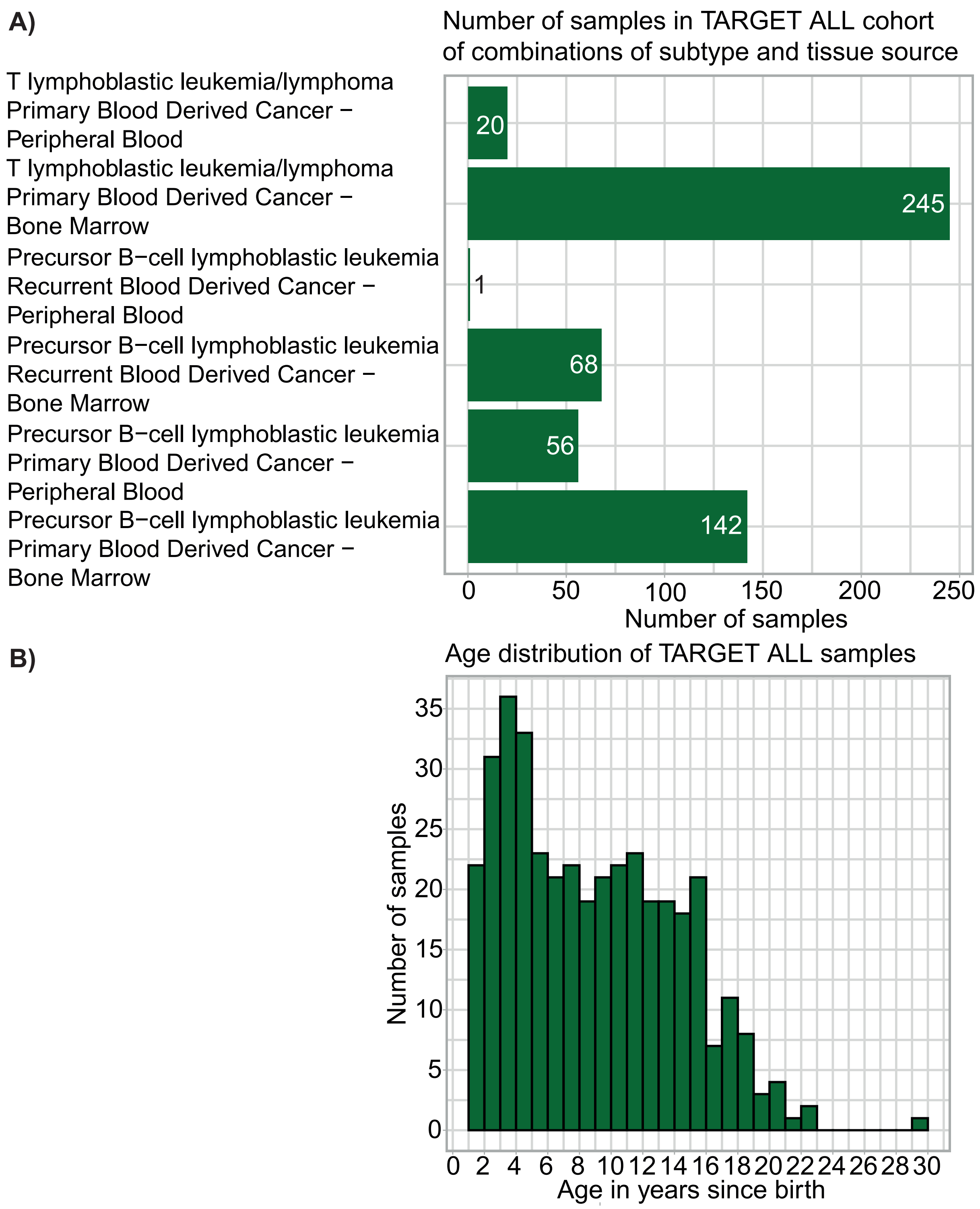


**Supplementary Figure S1.** Overview of the TARGET ALL cohort. **A)** Number of samples in the TARGET ALL cohort belonging to different combinations of tissue source, recurrence, and subtype. We retained only bone marrow primary samples for analyses as this category contained the largest group of samples and to ensure the direct comparison between the two subtypes was not confounded by differences in tissue type and recurrence. **B)** Distribution of age expressed in years since birth of TARGET ALL cohort subsetted to contain only bone marrow primary samples.
