## Supplementary Figure S2 for "Data-driven discovery of gene expression markers distinguishing pediatric acute lymphoblastic leukemia subtypes"

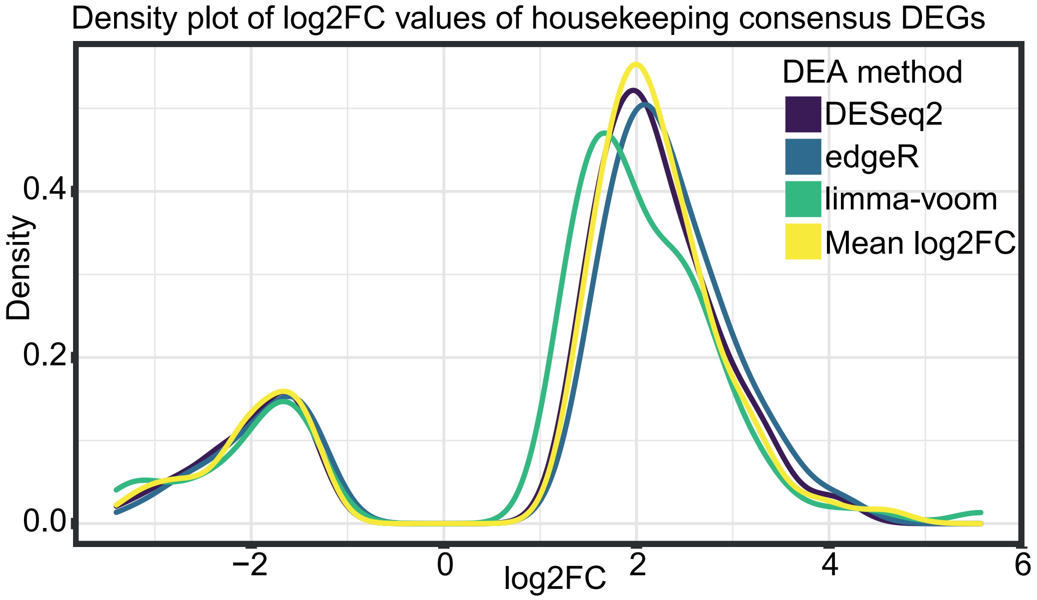


**Supplementary Figure S2.** Density plot of log2 fold change (log2FC) values of 103 housekeeping consensus differentially expressed genes (DEGs) predicted by three differential expression analysis (DEA) methods: limma-voom (green), edgeR (blue), and DESeq2 (purple). The distribution of the mean log2FC of the housekeeping DEGs across the three DEA methods is also shown (yellow). Housekeeping consensus DEGs were found by comparing the consensus DEGs with housekeeping genes reported by Eisenberg and Levanon (2013).
