## Supplementary Figure S3 for "Data-driven discovery of gene expression markers distinguishing pediatric acute lymphoblastic leukemia subtypes"

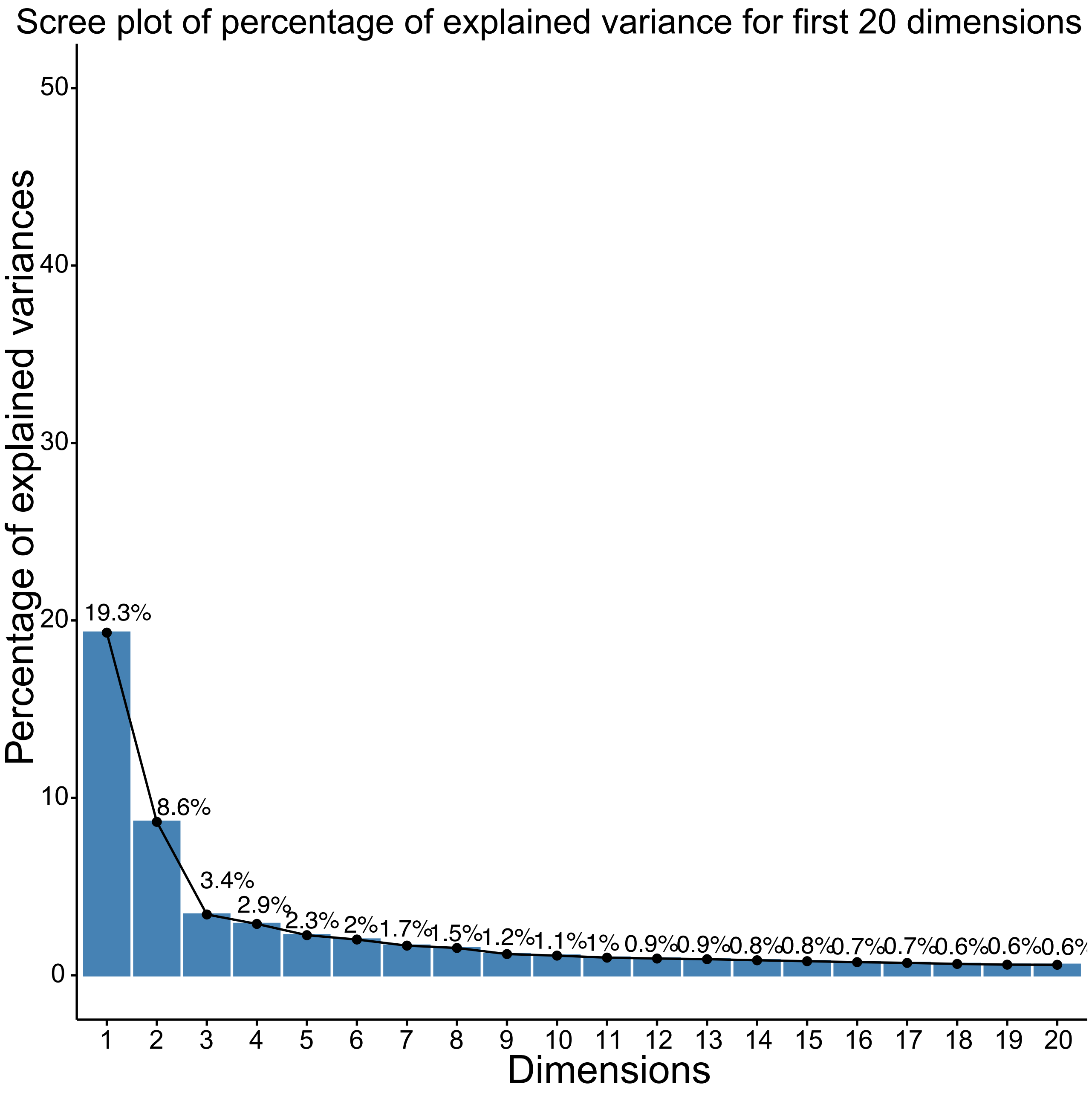


**Supplementary Figure S3.** Scree plot of percentage of explained variance for the first 20 principal component (PC) dimensions from principal component analysis (PCA). The percentage of explained variance for each of the first 20 dimensions are shown on top of each bar.
