## Supplementary Figure S4 for "Data-driven discovery of gene expression markers distinguishing pediatric acute lymphoblastic leukemia subtypes"

**A)**

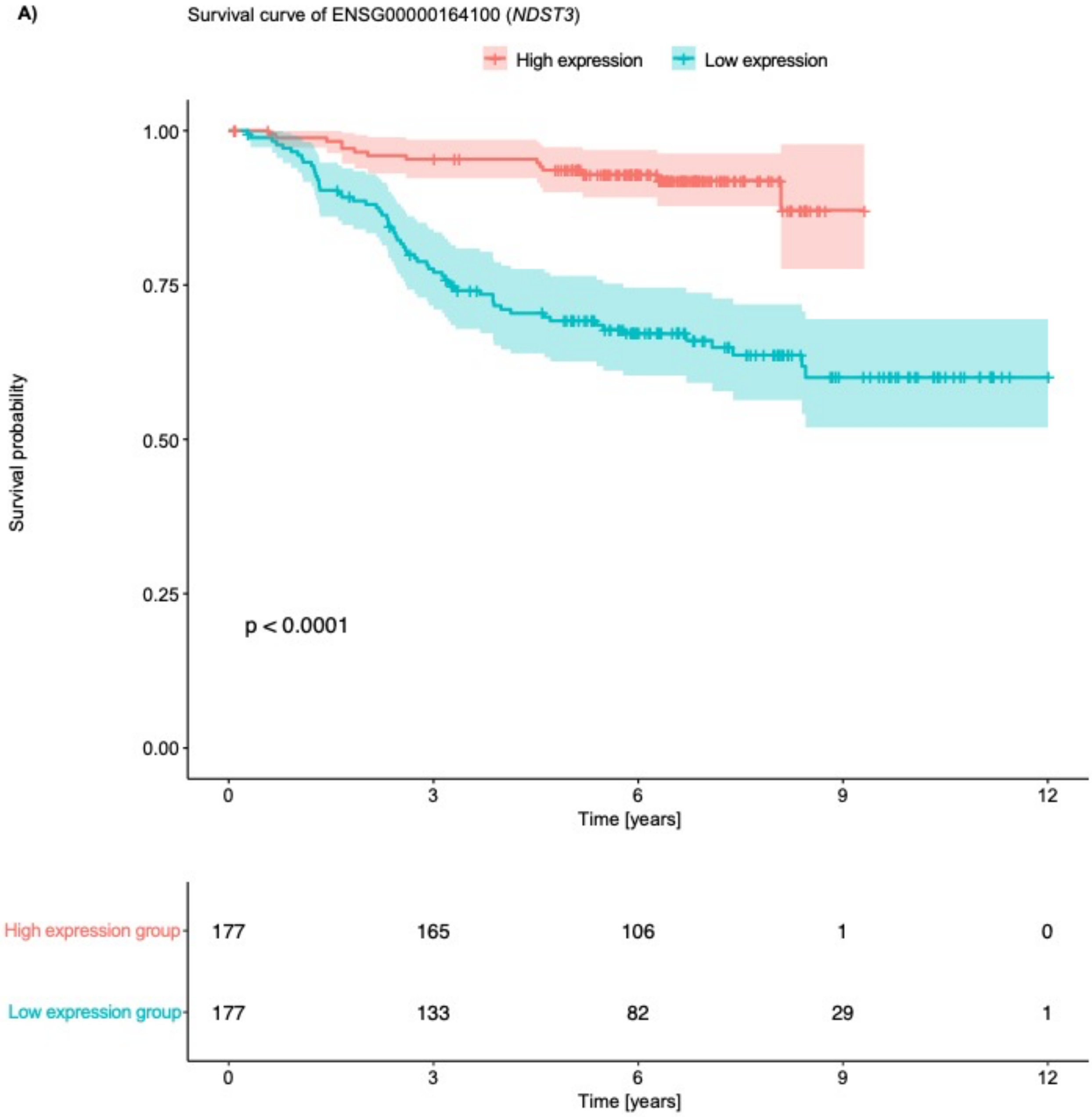
Survival curve of ENSG00000164100 (*NDST3*)
High expression
Low expression
Survival probability
1.00
0.75
0.50
0.25
0.00
p < 0.0001
0
3
6
9
12
Time [years]
High expression group
177
165
106
1
0
Low expression group
177
133
82
29
1
0
3
6
9
12
Time [years]

B)

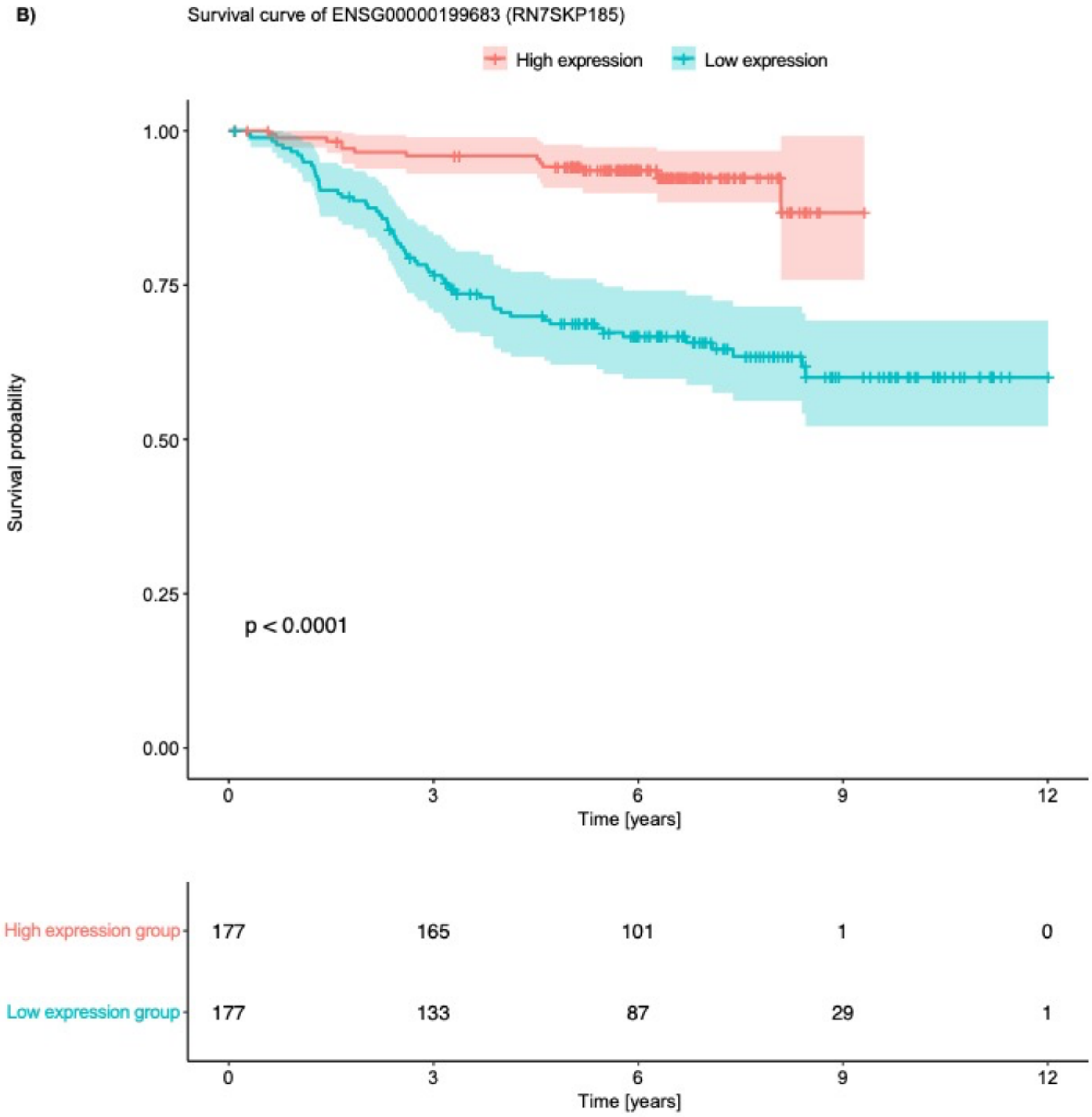

Survival curve of ENSG00000199683 (RN7SKP185)
High expression
Low expression
Survival probability
1.00
0.75
0.50
0.25
0.00
p < 0.0001
0
3
6
9
12
Time [years]
High expression group
177
165
101
1
0
Low expression group
177
133
87
29
1
0
3
6
9
12
Time [years]

C)

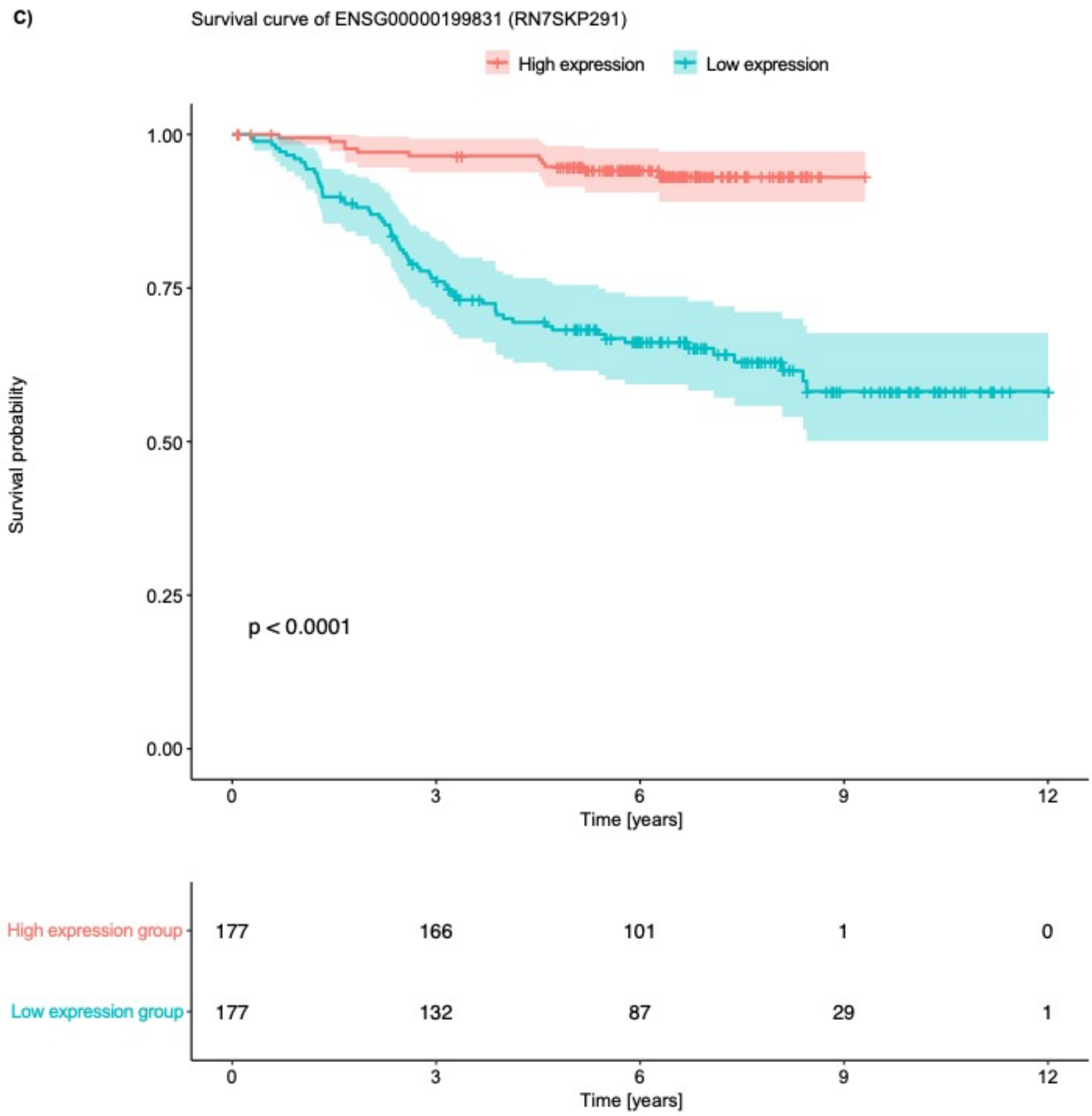

Survival curve of ENSG00000199831 (RN7SKP291)
High expression
Low expression
1.00
0.75
0.50
0.25
0.00
Survival probability
p < 0.0001
0
3
6
9
12
Time [years]
High expression group
Low expression group
177
166
101
1
0
177
132
87
29
1
0
3
6
9
12
Time [years]

D)

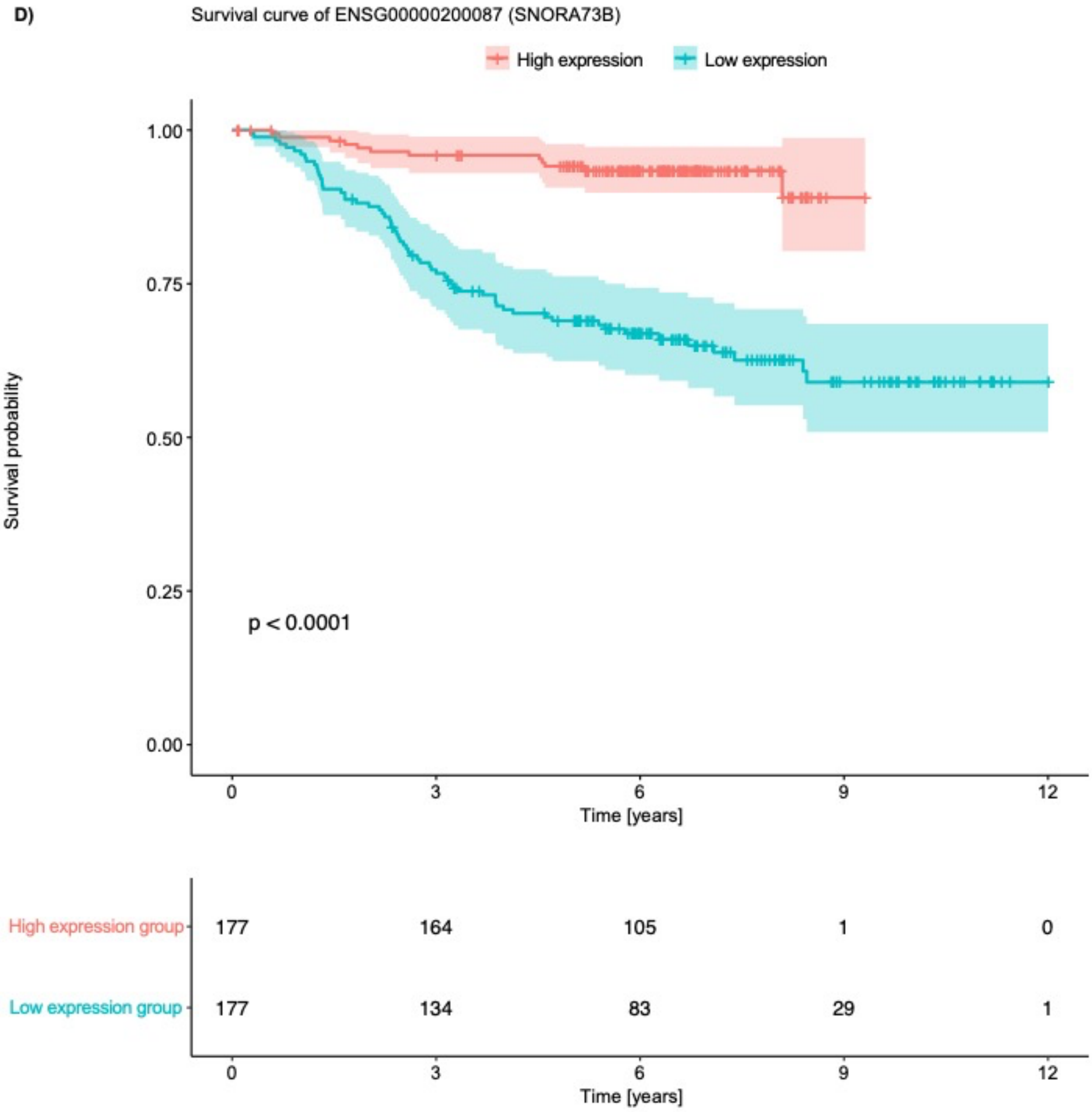
Survival curve of ENSG00000200087 (SNORA73B)
High expression
Low expression
Survival probability
1.00
0.75
0.50
0.25
0.00
p < 0.0001
0
3
6
9
12
Time [years]
High expression group
177
164
105
1
0
Low expression group
177
134
83
29
1
0
3
6
9
12
Time [years]

E)

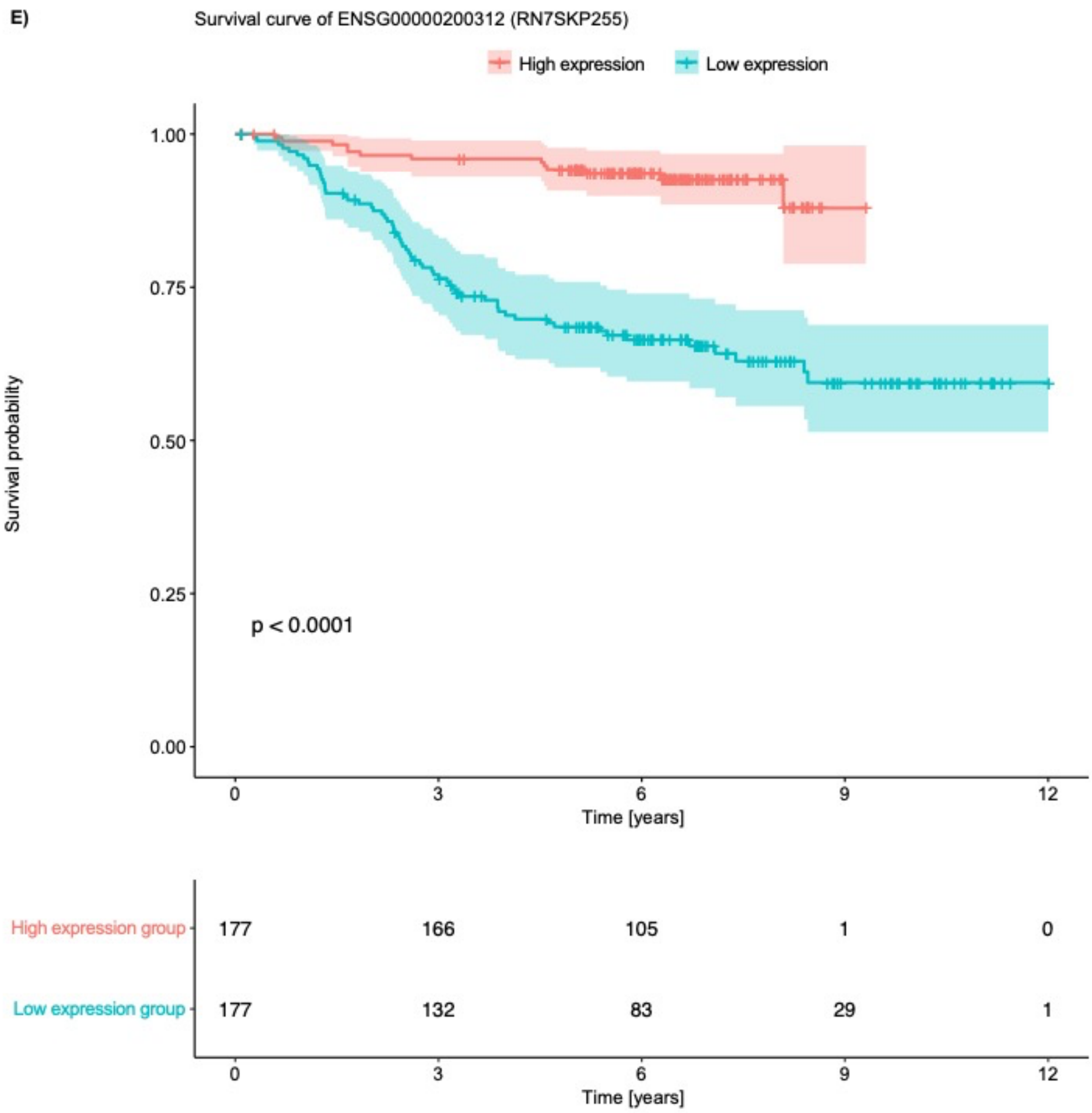

Survival curve of ENSG00000200312 (RN7SKP255)
High expression
Low expression
Survival probability
1.00
0.75
0.50
0.25
0.00
p < 0.0001
0
3
6
9
12
Time [years]
High expression group
Low expression group
177
166
105
1
0
177
132
83
29
1
0
3
6
9
12
Time [years]

F)

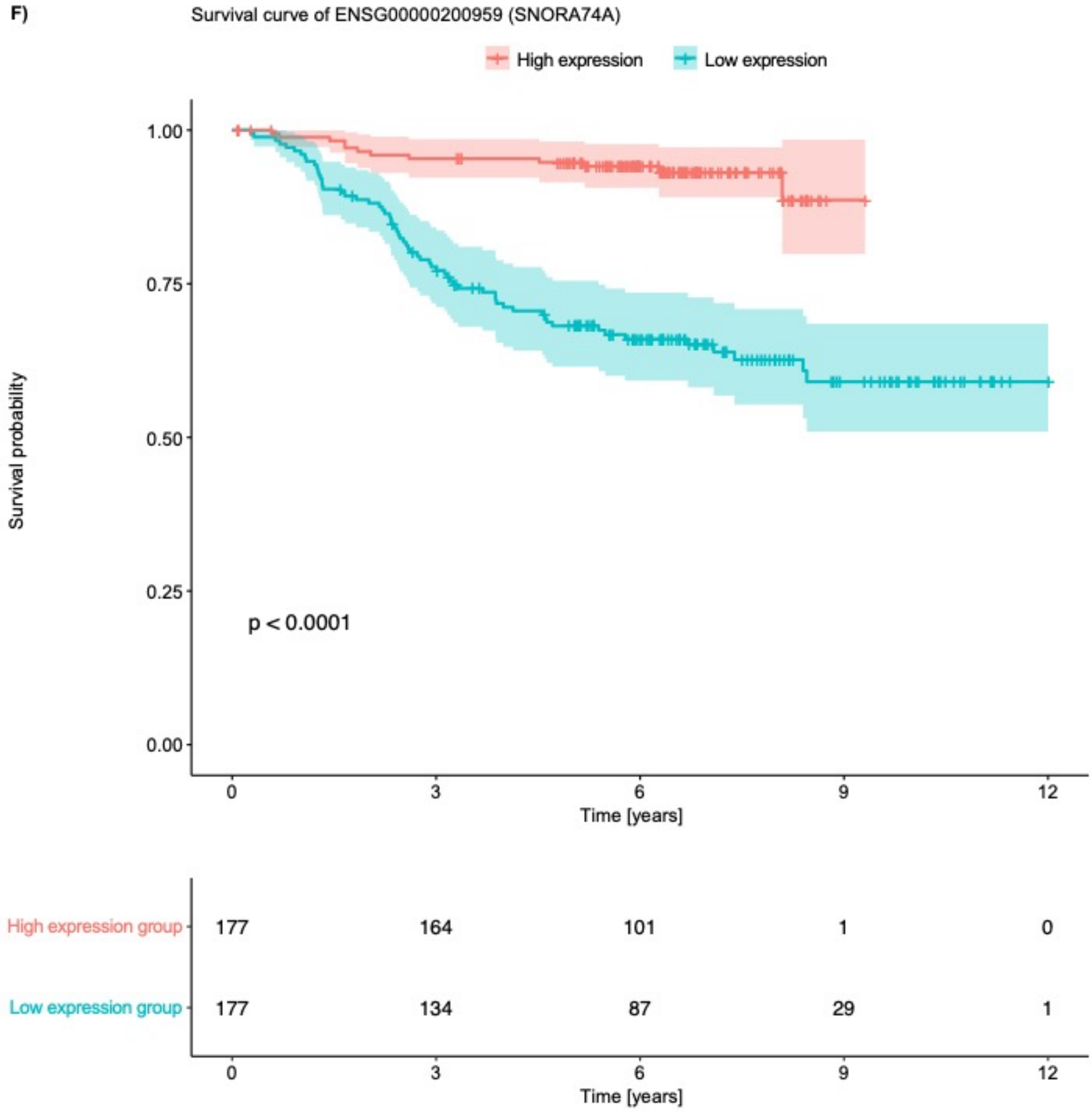
Survival curve of ENSG00000200959 (SNORA74A)
High expression
Low expression
Survival probability
1.00
0.75
0.50
0.25
0.00
p < 0.0001
0
3
6
9
12
Time [years]
High expression group
177
164
101
1
0
Low expression group
177
134
87
29
1
0
3
6
9
12
Time [years]

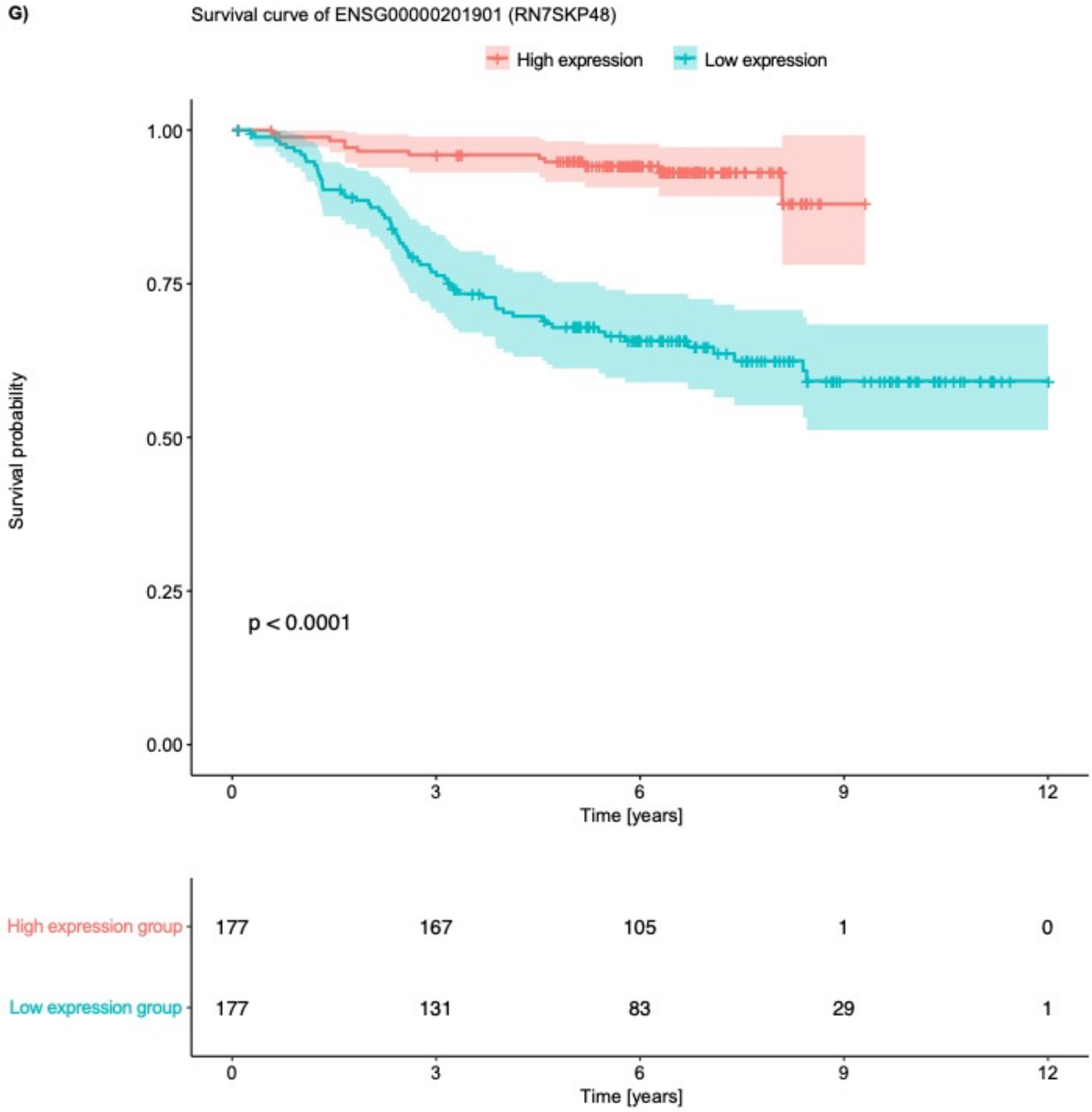
G)
Survival curve of ENSG00000201901 (RN7SKP48)
High expression
Low expression
Survival probability
1.00
0.75
0.50
0.25
0.00
p < 0.0001
0
3
6
9
12
Time [years]
High expression group
177
167
105
1
0
Low expression group
177
131
83
29
1
0
3
6
9
12
Time [years]

H)

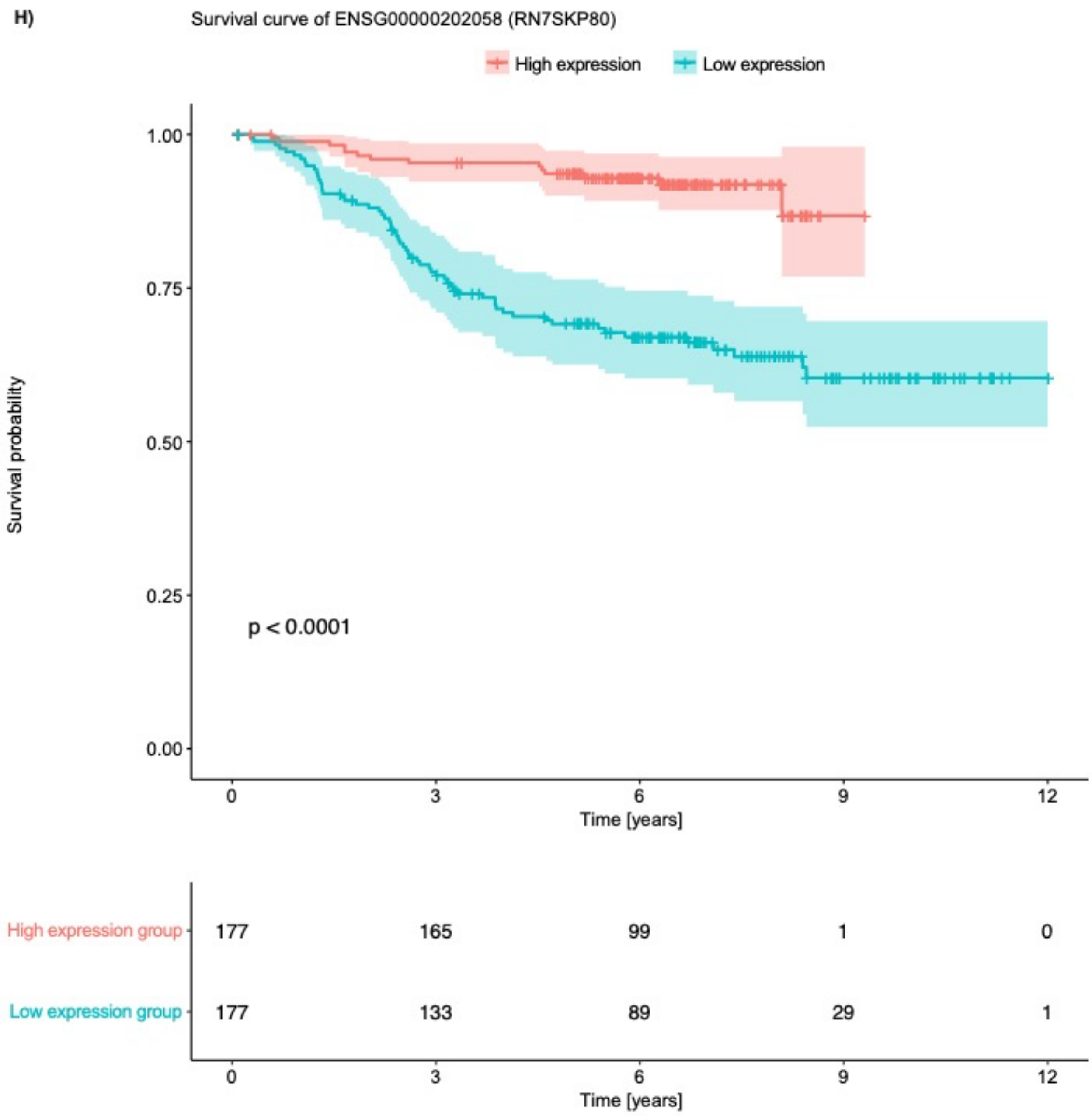
Survival curve of ENSG00000202058 (RN7SKP80)
High expression
Low expression
Survival probability
1.00
0.75
0.50
0.25
0.00
p < 0.0001
0
3
6
9
12
Time [years]
High expression group
177
165
99
1
0
Low expression group
177
133
89
29
1
0
3
6
9
12
Time [years]

**I)**

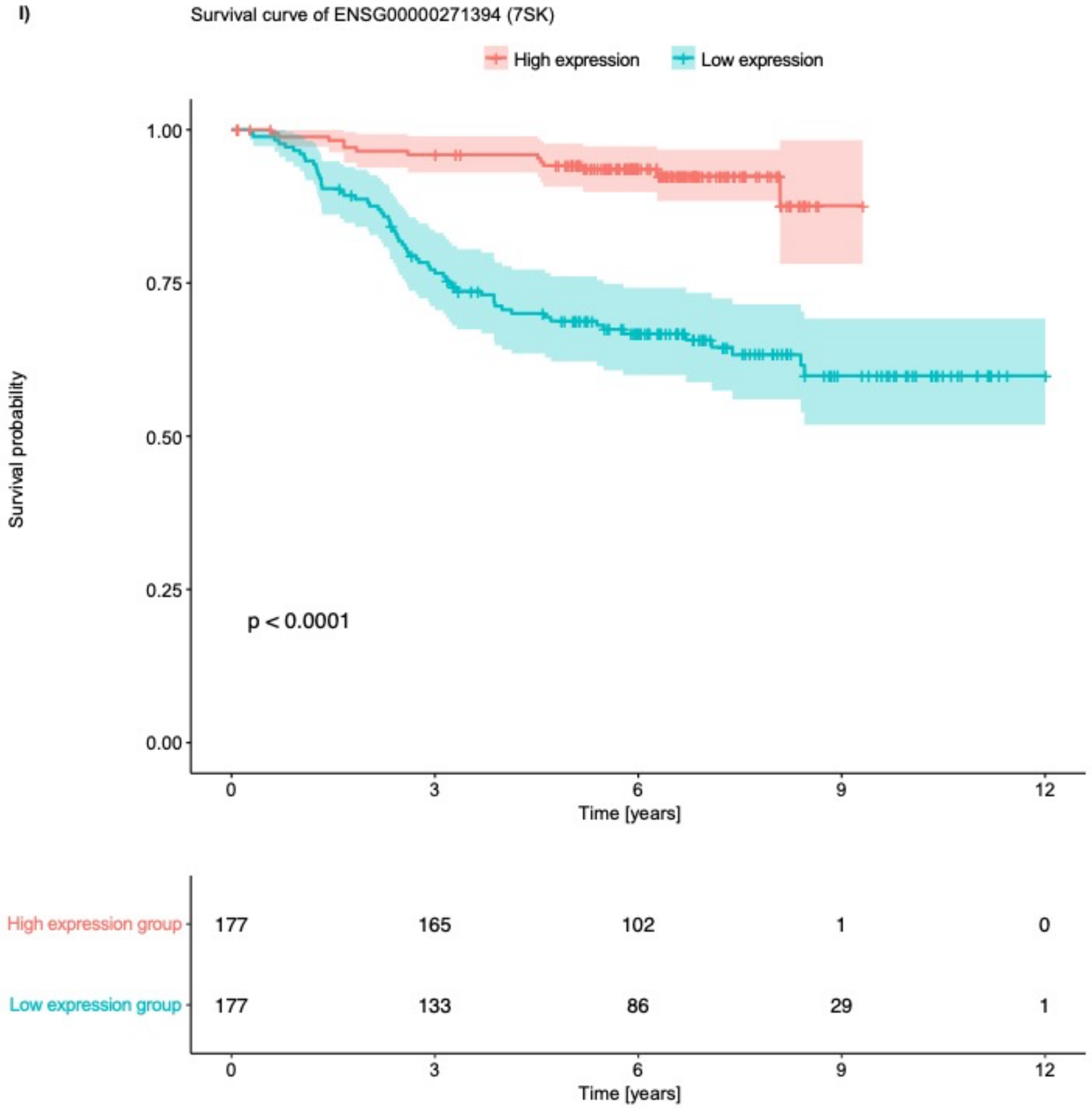


**Supplementary Figure S4.** Kaplan-Meier survival plots of nine of the discovered subtype-related gene expression markers: **A)** *NDST3*, **B)** RN7SKP185, **C)** RN7SKP291, **D)** SNORA73B, **E)** RN7SKP255, **F)** SNORA74A, **G)** RN7SKP48, **H)** RN7SKP80, and **I) 7SK.** Patients were categorized into high (orange) and low (blue) expression groups based on whether their expression values were above or below the median expression level of the corresponding gene. Survival curves were constructed using the discrete expression group as the independent variable, and the significance of the difference in survival between the two groups was assessed using a log-rank test with a *p*-value < 0.05 considered statistically significant.
