## Supplementary Figure S5 for "Data-driven discovery of gene expression markers distinguishing pediatric acute lymphoblastic leukemia subtypes"

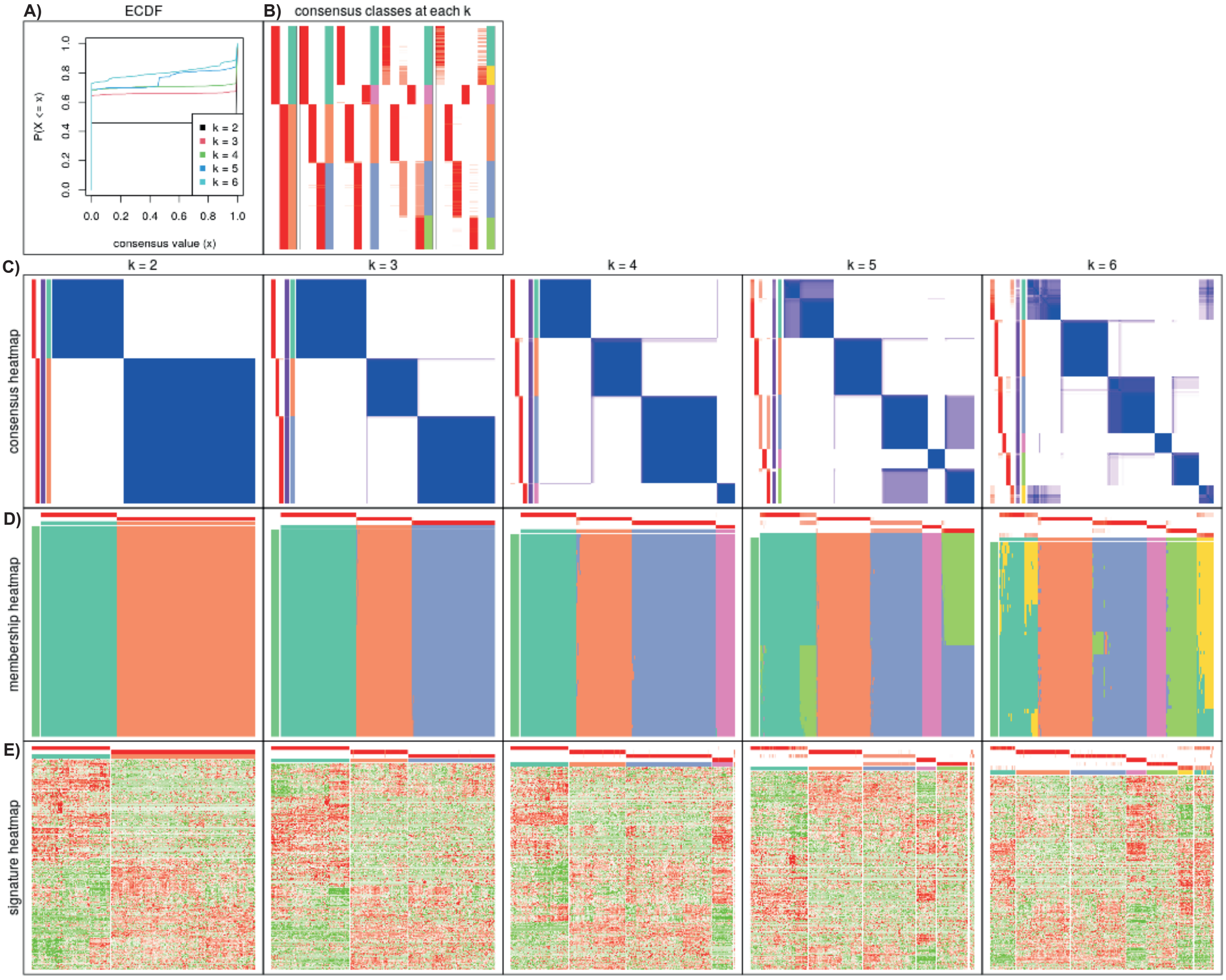


**Supplementary Figure S5.** Visualization of unsupervised clustering using method SD:mclust. The unsupervised clustering was performed using the *cola* framework. **A)** Empirical cumulative distribution function curve of the consensus matrix for all values of *k*. **B)** Heatmap of predicted classes showing probability of samples (rows) to belong to *k* number of classes. **C)** Consensus heatmaps showing probability of two samples to be in the same subgroup across all partitions. **D)** Membership heatmaps visualizing the subgroup label that each partition (rows) predicts the samples (columns) to belong to. **E)** Signature heatmaps showing the genes with significant differences in expression between subgroups.
