## Supplementary Figure S6 for "Data-driven discovery of gene expression markers distinguishing pediatric acute lymphoblastic leukemia subtypes"

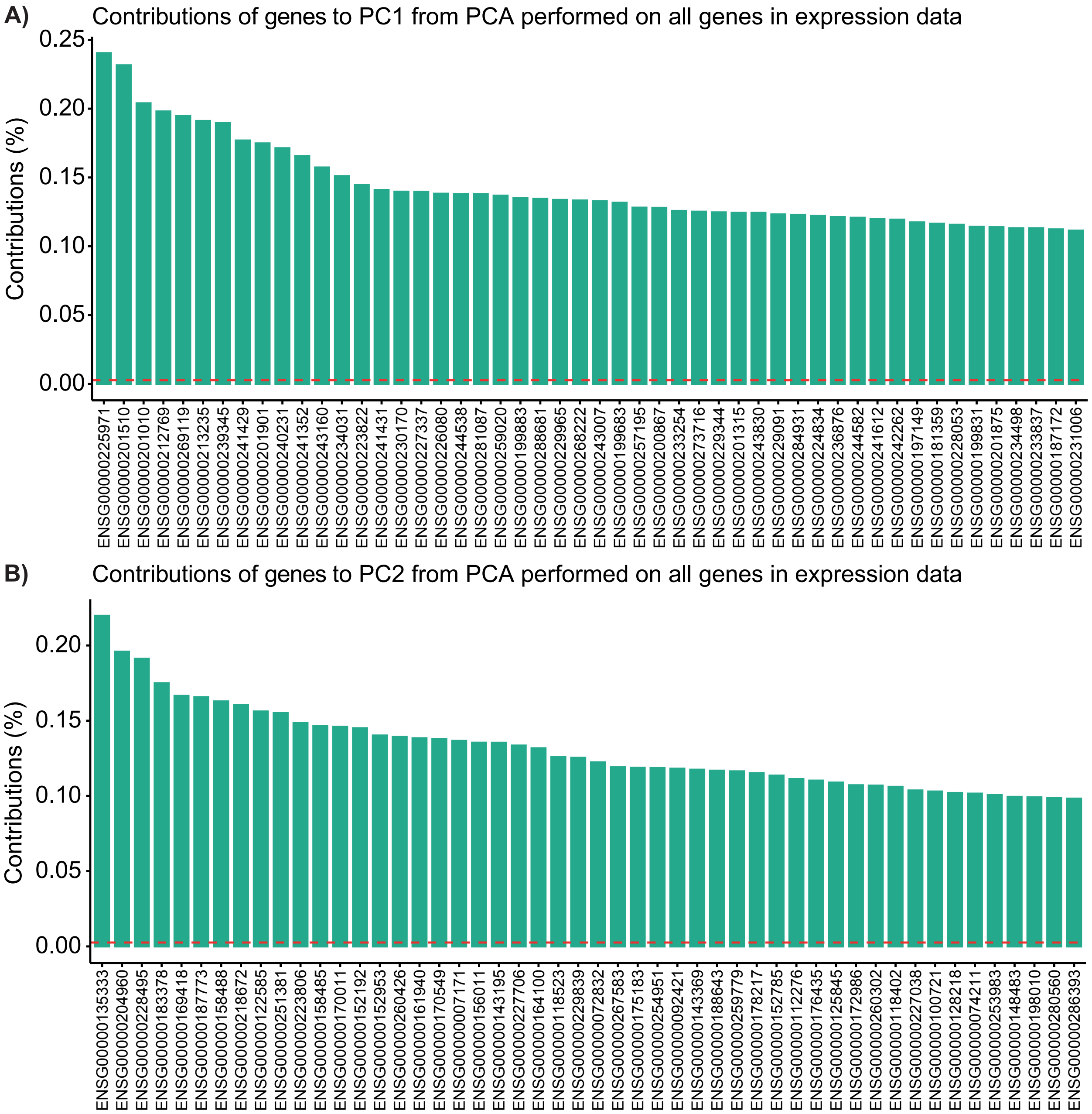


**Supplementary Figure S6.** Contributions in % of the top 50 genes to principal component 1 (PC1) **A)** and principal component 2 (PC2) **B)**. The principal component analysis (PCA) was performed on gene expression data of a Danish cohort of pediatric patients with ALL.
