## Supplementary Table S2 for "Data-driven discovery of gene expression markers distinguishing pediatric acute lymphoblastic leukemia subtypes"

| Supplementary Table S2. List of 103 housekeeping differentially expressed genes (DEGs). 103 DEGs overlapped between the list of housekeeping genes provided by Eisenberg and Levanon (2013) and our list of consensus DEGs. The average log2 fold change (log2FC) and average False Discovery Rate (FDR) values of the DEGs are shown (average across three differential expression analysis (DEA) methods: limma-voom, edgeR, and DESeq2). | | | |
|---|---|---|---|
| **ENSEMBL gene ID** | **External gene name** | **Average log2FC [SD]** | **Average FDR [SD]** |
| ENSG00000117528 | *ABCD3* | -2.0558 [0.2284] | 7.8268e-10 [1.3556e-09] |
| ENSG00000100439 | *ABHD4* | 2.7184 [0.2267] | 4.0769e-13 [7.0268e-13] |
| ENSG00000189046 | *ALKBH2* | -1.6685 [0.2218] | 1.1818e-03 [1.5969e-03] |
| ENSG00000197043 | *ANXA6* | -2.0312 [0.2928] | 6.7560e-08 [8.8843e-08] |
| ENSG00000132694 | *ARHGEF11* | 2.0676 [0.2478] | 1.9266e-03 [3.2068e-03] |
| ENSG00000136950 | *ARPC5L* | 2.1132 [0.4218] | 1.5462e-08 [2.6780e-08] |
| ENSG00000170653 | *ATF7* | 1.9854 [0.0941] | 3.4806e-24 [6.0287e-24] |
| ENSG00000017260 | *ATP2C1* | -2.0125 [0.2526] | 1.2952e-06 [2.2433e-06] |
| ENSG00000185883 | *ATP6V0C* | 3.1741 [0.7032] | 1.5037e-34 [2.6045e-34] |
| ENSG00000117475 | *BLZF1* | -1.7979 [0.1735] | 7.6949e-06 [1.3327e-05] |

| | | | |
|---|---|---|---|
| ENSG00000141867 | *BRD4* | 1.9672 [0.1583] | 3.4038e-08 [5.6954e-08] |
| ENSG00000171067 | *C11orf24* | 1.9540 [0.3260] | 2.1345e-04 [3.6970e-04] |
| ENSG00000152492 | *CCDC50* | 4.5889 [0.8558] | 3.0237e-27 [5.1813e-27] |
| ENSG00000111666 | *CHPT1* | 1.7283 [0.1110] | 2.2142e-04 [3.8067e-04] |
| ENSG00000140848 | *CPNE2* | 2.7974 [0.3735] | 4.5742e-18 [7.92270e-18] |
| ENSG00000160741 | *CRTC2* | 2.2752 [0.4505] | 7.6463e-07 [1.3244e-06] |
| ENSG00000141551 | *CSNK1D* | 1.7483 [0.3589] | 1.1075e-02 [1.9182e-02] |
| ENSG00000177613 | *CSTF2T* | -1.3646 [0.0220] | 3.2144e-03 [4.3306e-03] |
| ENSG00000044115 | *CTNNA1* | 2.2630 [0.0640] | 2.3263e-11 [4.0158e-11] |
| ENSG00000055130 | *CUL1* | 2.0240 [0.1984] | 2.3276e-07 [4.0132e-07] |
| ENSG00000100243 | *CYB5R3* | 1.5671 [0.2937] | 1.2558e-02 [2.1751e-02] |
| ENSG00000273749 | *CYFIP1* | 2.5279 [0.1503] | 4.3537e-14 [7.5153e-14] |
| ENSG00000155016 | *CYP2U1* | -3.2764 [0.0745] | 3.9247e-10 [6.797767e-10] |

| | | | |
|---|---|---|---|
| ENSG00000164535 | *DAGLB* | 3.0061 [0.2123] | 4.6799e-32 [8.1057e-32] |
| ENSG00000112679 | *DUSP22* | 1.3907 [0.0851] | 1.8042e-02 [2.2279e-02] |
| ENSG00000146425 | *DYNLT1* | 1.5265 [0.0763] | 3.3111e-03 [2.8786e-03] |
| ENSG00000173812 | *EIF1* | 1.8970 [0.5505] | 9.1656e-03 [1.5875e-02] |
| ENSG00000114784 | *EIF1B* | 1.9802 [0.2660] | 1.4193e-06 [2.4583e-06] |
| ENSG00000172071 | *EIF2AK3* | 2.6319 [0.0589] | 1.0763e-18 [1.8640e-18] |
| ENSG00000118985 | *ELL2* | 2.8740 [0.1208] | 1.8080e-13 [1.5438e-13] |
| ENSG00000063245 | *EPN1* | 1.9252 [0.2762] | 1.4290e-12 [2.4751e-12] |
| ENSG00000137166 | *FOXP4* | 2.3874 [0.6924] | 4.0462e-04 [7.0081e-04] |
| ENSG00000136877 | *FPGS* | 2.0930 [0.6648] | 1.5153e-03 [2.6245e-03] |
| ENSG00000142252 | *GEMIN7* | 2.0477 [0.2924] | 4.1136e-09 [7.1250e-09] |
| ENSG00000113552 | *GNPDA1* | -2.6470 [0.0455] | 3.9711e-09 [6.8768e-09] |
| ENSG00000114745 | *GORASP1* | 1.5397 [0.2276] | 2.3025e-03 [3.9881e-03] |

| | | | |
|---|---|---|---|
| ENSG00000013583 | *HEBP1* | 2.1593 [0.0889] | 2.5504e-07 [3.5537e-07] |
| ENSG00000131724 | *IL13RA1* | 2.1791 [0.2429] | 7.8104e-04 [1.2863e-03] |
| ENSG00000205730 | *ITPRIPL2* | 4.0302 [0.0750] | 2.6791e-20 [4.6404e-20] |
| ENSG00000109787 | *KLF3* | 2.1830 [0.2595] | 2.0987e-05 [3.5030e-05] |
| ENSG00000119138 | *KLF9* | 2.6559 [0.1379] | 5.7215e-10 [4.0431e-10] |
| ENSG00000135686 | *KLHL36* | 1.9373 [0.0346] | 1.5489e-10 [1.7241e-10] |
| ENSG00000163155 | *LYSMD1* | -1.4843 [0.0323] | 7.4410e-03 [1.0130e-02] |
| ENSG00000068305 | *MEF2A* | 1.4044 [0.0526] | 6.4894e-03 [1.0999e-02] |
| ENSG00000100139 | *MICALL1* | 1.9195 [0.5175] | 6.1831e-03 [1.0709e-02] |
| ENSG00000175806 | *MSRA* | 3.7525 [0.6172] | 1.1157e-20 [1.7109e-20] |
| ENSG00000170873 | *MTSS1* | 1.6227 [0.0843] | 1.0680e-02 [8.7850e-03] |
| ENSG00000188554 | *NBR1* | -1.6572 [0.0464] | 3.3377e-08 [5.21981e-08] |
| ENSG00000070614 | *NDST1* | 2.5957 [0.0510] | 3.7965e-10 [4.6511e-10] |

| | | | |
|---|---|---|---|
| ENSG00000141458 | *NPC1* | 1.9508 [0.2024] | 9.5397e-08 [1.6523e-07] |
| ENSG00000119655 | *NPC2* | 2.4175 [0.3626] | 3.2154e-07 [5.5692e-07] |
| ENSG00000103148 | *NPRL3* | 1.6068 [0.1496] | 2.1433e-07 [3.7100e-07] |
| ENSG00000124588 | *NQO2* | -1.7752 [0.2779 | 1.2586e-05 [1.9585e-05] |
| ENSG00000151623 | *NR3C2* | 2.4396 [0.5887] | 7.7398e-03 [1.3406e-02] |
| ENSG00000157045 | *NTAN1* | 1.6784 [0.1135] | 9.2072e-08 [1.5895e-07] |
| ENSG00000183828 | *NUDT14* | 3.1752 [0.1034] | 3.1144e-18 [5.3942e-18] |
| ENSG00000217930 | *PAM16* | 1.6526 [0.2599] | 1.6941e-04 [2.9343e-04] |
| ENSG00000163110 | *PDLIM5* | -2.2709 [0.1124] | 3.9527e-07 [6.8447e-07] |
| ENSG00000152684 | *PELO* | -2.8975 [0.2320] | 1.2503e-16 [1.1482e-16] |
| ENSG00000166821 | *PEX11A* | -3.0641 [0.3066] | 3.7929e-13 [6.5656e-13] |
| ENSG00000139197 | *PEX5* | -2.6268 [0.3444] | 4.0208e-17 [6.9643e-17] |
| ENSG00000130517 | *PGPEP1* | -2.1281 [0.4174] | 2.4834e-07 [2.6502e-07] |

| | | | |
|---|---|---|---|
| ENSG00000272391 | *POM121C* | 1.6201 [0.0851] | 1.7061e-07 [2.9195e-07] |
| ENSG00000112033 | *PPARD* | 2.4167 [0.3865] | 1.2427e-11 [2.1524e-11] |
| ENSG00000143801 | *PSEN2* | 2.3613 [0.2920] | 7.8223e-04 [1.3541e-03] |
| ENSG00000130348 | *QRSL1* | 1.8480 [0.1013] | 2.8844e-09 [3.9329e-09] |
| ENSG00000137040 | *RANBP6* | -1.5199 [0.1505] | 2.2870e-03 [3.9608e-03] |
| ENSG00000107263 | *RAPGEF1* | 1.7829 [0.2424] | 1.7698e-05 [3.0654e-05] |
| ENSG00000183808 | *RBM12B* | -1.4516 [0.0764] | 2.4313e-03 [4.1415e-03] |
| ENSG00000159200 | *RCAN1* | 2.3074 [0.1182] | 2.7208e-06 [4.1817e-06] |
| ENSG00000129625 | *REEP5* | 1.7002 [0.1148] | 1.6964e-05 [2.9292e-05] |
| ENSG00000173039 | *RELA* | 1.9231 [0.5209] | 3.5427e-04 [6.1361e-04] |
| ENSG00000143878 | *RHOB* | 3.3279 [0.3322] | 5.6136e-12 [9.7224e-12] |
| ENSG00000132669 | *RIN2* | 3.3818 [0.0562] | 6.1996e-20 [6.8905e-20] |
| ENSG00000151748 | *SAV1* | 2.1427 [0.4001] | 4.1695e-07 [7.1196e-07] |

| | | | |
|---|---|---|---|
| ENSG00000138760 | *SCARB2* | 1.8510 [0.3696] | 1.3454e-03 [2.2869e-03] |
| ENSG00000124570 | *SERPINB6* | 2.5993 [0.5928] | 4.5298e-05 [6.7008e-05] |
| ENSG00000102100 | *SLC35A2* | 1.9292 [0.2430] | 1.4498e-06 [2.5111e-06] |
| ENSG00000175782 | *SLC35E3* | 2.6622 [0.0783] | 4.7145e-26 [8.1658e-26] |
| ENSG00000205302 | *SNX2* | 2.5882 [0.2717] | 3.7825e-38 [3.4408e-38] |
| ENSG00000109762 | *SNX25* | 2.2784 [0.1487] | 2.6136e-05 [3.6068e-05] |
| ENSG00000130340 | *SNX9* | 3.0137 [0.0850] | 5.2106e-13 [7.8938e-13] |
| ENSG00000123178 | *SPRYD7* | 2.0087 [0.0868] | 6.9119e-08 [5.9435e-08] |
| ENSG00000100104 | *SRRD* | 1.3270 [0.0526] | 4.6980e-04 [4.2679e-04] |
| ENSG00000040341 | *STAU2* | -2.1003 [0.1069] | 2.9640e-09 [5.1337e-09] |
| ENSG00000100242 | *SUN2* | 1.5278 [0.1522] | 2.1608e-03 [3.7423e-03] |
| ENSG00000162298 | *SYVN1* | 3.0607 [0.3542] | 3.1243e-45 [5.4115e-45] |
| ENSG00000169762 | *TAPT1* | 1.5817 [0.0339] | 1.6043e-03 [2.4792e-03] |

| | | | |
|---|---|---|---|
| ENSG00000132604 | *TERF2* | 2.0445 [0.2279] | 6.1102e-11 [1.0583e-10] |
| ENSG00000127666 | *TICAM1* | 2.3204 [0.4467] | 4.8051e-08 [8.3228e-08] |
| ENSG00000196781 | *TLE1* | 3.1004 [0.0917] | 7.3106e-14 [1.2418e-13] |
| ENSG00000137842 | *TMEM62* | 2.0251 [0.3619] | 3.8211e-06 [6.5942e-06] |
| ENSG00000180694 | *TMEM64* | 2.4424 [0.8092] | 5.6949e-04 [9.2631e-04] |
| ENSG00000125827 | *TMX4* | 1.6811 [0.1878] | 8.2547e-05 [1.4190e-04] |
| ENSG00000243725 | *TTC4* | -1.4174 [0.1327] | 6.5920e-03 [6.6144e-03] |
| ENSG00000153443 | *UBALD1* | 2.5131 [0.5384] | 2.8853e-16 [4.9974e-16] |
| ENSG00000198833 | *UBE2J1* | 1.5392 [0.1807] | 4.2615e-03 [7.2222e-03] |
| ENSG00000115652 | *UXS1* | 1.3878 [0.1048] | 4.5820e-03 [7.8179e-03] |
| ENSG00000140006 | *WDR89* | -1.7109 [0.0459] | 5.5711e-09 [6.4233e-09] |
| ENSG00000175155 | *YPEL2* | 2.0539 [0.1143] | 3.7284e-10 [6.3663e-10] |
| ENSG00000147905 | *ZCCHC7* | 2.5712 [0.1514] | 1.2016e-40 [2.0804e-40] |

| | | | |
|---|---|---|---|
| ENSG00000178764 | *ZHX2* | 2.1086 [0.2899] | 1.1163e-10 [1.9334e-10] |
| ENSG00000179195 | *ZNF664* | -1.4443 [0.1614] | 2.7303e-03 [3.8936e-03] |

Abbreviations: DEGs, differentially expressed genes; log2FC, log2 fold change; FDR, false discovery rate; DEA, differential expression analysis; SD, standard deviation.
