## Supplementary Table S3 for "Data-driven discovery of gene expression markers distinguishing pediatric acute lymphoblastic leukemia subtypes"

**Supplementary Table S3. Consensus set of selected ENSEMBL gene IDs from elastic net binomial logistic regression together with their external gene name and average coefficient sorted in descending order by absolute average coefficient. We created the consensus set of ENSEMBL gene IDs based on overlap of selected ENSEMBL gene IDs from 10 elastic net logistic regression runs. The consensus genes are compared with results of the consensus differential expression analysis (DEA).**

| ENSEMBL gene ID | External gene name [biotype] | Average elastic net coefficient [SD] | Average log2FC [SD] | Average FDR [SD] |
|---|---|---|---|---|
| ENSG00000243062 | Novel gene [lncRNA] | 0.0179 [0.002] | -5.4104 [0.5948] | 7.0414e-53 [1.1355e-52] |
| ENSG00000199831 | RN7SKP291 [misc RNA] | 0.0137 [0.0004] | -9.3511 [0.2172] | 5.5305e-87 [9.5791e-87] |
| ENSG00000202058 | RN7SKP80 [misc RNA] | 0.0116 [0.0003] | -8.7568 [0.7806] | 3.3216e-80 [5.7532e-80] |
| ENSG00000228997 | Novel gene [processed pseudogene] | 0.0087 [0.0018] | -5.6471 [0.1568] | 3.7650e-42 [6.5211e-42] |
| ENSG00000275155 | Novel gene [lncRNA] | -0.0080 [0.0013] | 6.4042 [0.3272] | 3.8950e-79 [6.7463e-79] |
| ENSG00000202357 | Y_RNA [misc RNA] | 0.0079 [0.0009] | -7.9692 [0.7869] | 3.6584e-55 [6.3366e-55] |
| ENSG00000164100 | NDST3 [protein coding] | 0.0076 [0.0005] | -11.6426 [0.3028] | 2.1479e-77 [3.7186e-77] |
| ENSG00000199683 | RN7SKP185 [misc RNA] | 0.0073 [0.0007] | -7.7758 [0.6542] | 1.1201e-70 [1.5208e-70] |
| ENSG00000279204 | Novel gene [TEC] | -0.0067 [0.0007] | 7.2987 [0.1192] | 6.1784e-92 [1.0701e-91] |
| ENSG00000095585 | BLNK [protein coding] | -0.0063 [0.0009] | 7.0636 [0.4600] | 7.1782e-93 [1.2433e-92] |
| ENSG00000223806 | LINC00114 [lncRNA] | -0.0062 [0.0009] | 8.7047 [0.2038] | 2.9924e-87 [5.1830e-87] |
| ENSG00000225206 | MIR137HG [lncRNA] | 0.0055 [0.0006] | -7.5747 [0.5870] | 3.1520e-52 [5.4594e-52] |
| ENSG00000118523 | CCN2 [protein coding] | -0.0054 [0.0010] | 9.3982 [0.2464] | 2.3999e-81 [4.1568e-81] |

| | | | | |
|---|---|---|---|---|
| ENSG00000238741 | SCARNA7 [scaRNA] | 0.0054 [0.0011] | -6.7528 [0.0907] | 1.8558e-48 [3.2144e-48] |
| ENSG00000200959 | SNORA74A [snoRNA] | 0.0052 [0.0006] | -8.6670 [0.3967] | 1.3186e-58 [2.2837e-58] |
| ENSG00000200087 | SNORA73B [snoRNA] | 0.0050 [0.0005] | -7.3992 [0.9972] | 2.1027e-44 [3.6421e-44] |
| ENSG00000263413 | MIR4538 [miRNA] | -0.0049 [0.0024] | 6.6022 [1.6395] | 1.8674e-66 [3.2344e-66] |
| ENSG00000105694 | ELOCP28 [processed pseudogene] | -0.0047 [0.0022] | 6.3175 [0.1198] | 6.6892e-59 [1.1586e-58] |
| ENSG00000201901 | RN7SKP48 [misc RNA] | 0.0042 [0.0005] | -7.6915 [1.0032] | 2.3876e-47 [4.1355e-47] |
| ENSG00000177455 | CD19 [protein coding] | -0.0042 [0.0009] | 7.4110 [0.1468] | 2.3812e-118 [4.1243e-118] |
| ENSG00000212232 | SNORD17 [snoRNA] | 0.0041 [0.0009] | -7.0919 [0.6947] | 5.0064e-42 [8.6714e-42] |
| ENSG00000164330 | EBF1 [protein coding] | -0.0038 [0.0008] | 8.1850 [0.7341] | 9.4079e-103 [1.6295e-102] |
| ENSG00000200795 | RNU4-1 [snRNA] | 0.0035 [0.0007] | -6.6895 [0.4326] | 8.4889e-45 [1.4703e-44] |
| ENSG00000196092 | PAX5 [protein coding] | -0.0033 [0.0007] | 7.5755 [0.2704] | 3.9762e-107 [6.8870e-107] |
| ENSG00000200312 | RN7SKP255 [misc RNA] | 0.0029 [0.0006] | -8.1543 [1.5475] | 8.8446e-38 [1.5319e-37] |
| ENSG00000145642 | SHISAL2B [protein coding] | 0.0029 [0.0014] | -5.3578 [0.5424] | 2.1131e-51 [3.6599e-51] |
| ENSG00000183918 | SH2D1A [protein coding] | 0.0027 [0.0016] | -5.7853 [0.0451] | 4.6729e-49 [8.0937e-49] |
| ENSG00000227706 | Novel gene [lncRNA] | -0.0026 [0.0009] | 11.6906 [1.7298] | 1.0997e-90 [1.9047e-90] |
| ENSG00000200314 | Y_RNA [misc RNA] | 0.0017 [0.0013] | -8.3174 [0.8292] | 1.5658e-65 [2.7120e-65] |
| ENSG00000271394 | 7SK [misc RNA] | 0.0014 [0.0007] | -7.4783 [0.7991] | 3.7501e-47 [6.4834e-47] |
| ENSG00000128218 | VPREB3 [protein coding] | -0.0011 [0.0007] | 9.2479 [0.4986] | 1.0546e-106 [1.8266e-106] |

Abbreviations: DEA, differential expression analysis; SD, standard deviation; log2FC, log2 fold change; FDR, false discovery rate.
