## Supplementary Table S4 for "Data-driven discovery of gene expression markers distinguishing pediatric acute lymphoblastic leukemia subtypes"

| **Supplementary Table S4. Top 40 ENSEMBL gene IDs with highest contribution of explained variance between acute lymphoblastic leukemia (ALL) samples along the first principal component. ENSEMBL gene ID, external gene, biotype, and contribution of these 40 ENSEMBL gene IDs sorted by contribution in percentage. The ENSEMBL gene IDs are compared with results of the consensus differential expression analysis (DEA).** | | | | |
|---|---|---|---|---|
| **ENSEMBL gene ID** | **External gene name [biotype]** | **Contribution in %** | **Average log2FC [SD]** | **Average FDR [SD]** |
| ENSG00000164100 | NDST3 [protein coding] | 0.171 | -11.6426 [0.3028] | 2.1470e-77 [3.7186e-77] |
| ENSG00000158488 | CD1E [protein coding] | 0.1444 | -11.4757 [1.0199] | 7.1443e-16 [1.2374e-15] |
| ENSG00000158485 | CD1B [protein coding] | 0.1349 | -10.5611 [0.6251] | 5.8946e-18 [1.0210e-17] |
| ENSG00000200312 | RN7SKP255 [miscRNA] | 0.1252 | -8.1543 [1.5475] | 8.8446e-38 [1.5319e-37] |
| ENSG00000200488 | RN7SKP203 [miscRNA] | 0.1173 | -7.9260 [1.4508] | 1.6326e-36 [2.8277e-36] |
| ENSG00000227706 | Novel gene [lncRNA] | 0.1136 | 11.6906 [1.7298] | 1.0997e-90 [1.9047e-90] |
| ENSG00000202058 | RN7SKP80 [miscRNA] | 0.1099 | -8.7568 [0.7806] | 3.3216e-80 [5.7532e-80] |
| ENSG00000199831 | RN7SKP291 [miscRNA] | 0.107 | -9.3511 [0.2172] | 5.5305e-87 [9.5791e-87] |
| ENSG00000204960 | BLACE [lncRNA] | 0.1023 | 10.4528 [0.9408] | 1.2670e-59 [2.1946e-59] |
| ENSG00000118523 | CCN2 [protein coding] | 0.1011 | 9.3982 [0.2464] | 2.3999e-81 [4.1568e-81] |
| ENSG00000118402 | ELOVL4 [protein coding] | 0.0999 | -9.0693 [0.2217] | 3.1359e-19 [5.4316e-19] |
| ENSG00000172986 | GXYLT2 [protein coding] | 0.0977 | -7.4779 [1.2998] | 2.1511e-30 [2.4827e-30] |
| ENSG00000138650 | PCDH10 [protein coding] | 0.0973 | -8.0614 [0.9515] | 2.5417e-16 [4.3634e-16] |
| ENSG00000196581 | AJAP1 [protein coding] | 0.0961 | -5.7711 [2.4977] | 7.3267e-09 [1.2690e-08] |
| ENSG00000201 | RN7SKP48 | 0.0952 | -7.6915 [1.0032] | 2.3876e-47 |

| | | | | |
|---|---|---|---|---|
| 901 | [miscRNA] | | | [4.1355e-47] |
| ENSG00000228495 | LINC01013 [lncRNA] | 0.0948 | 8.6272 [0.2294] | 4.3675e-89 [7.5647e-89] |
| ENSG00000164330 | EBF1 [protein coding] | 0.0941 | 8.1850 [0.7341] | 9.4078e-103 [1.6295e-102] |
| ENSG00000236656 | Novel gene [lncRNA] | 0.094 | -10.4154 [2.1106] | 3.6629e-16 [6.3444e-16] |
| ENSG00000200959 | SNORA74A [snoRNA] | 0.0938 | -8.6670 [0.3967] | 1.3186e-58 [2.2837e-58] |
| ENSG00000223806 | LINC00114 [lncRNA] | 0.0927 | 8.7047 [0.2038] | 2.9924e-87 [5.1830e-87] |
| ENSG00000271394 | 7SK [miscRNA] | 0.0914 | -7.4783 [0.7991] | 3.7501e-47 [6.4834e-47] |
| ENSG00000100721 | TCL1A [protein coding] | 0.0896 | 8.8481 [0.1984] | 5.3461e-85 [9.2597e-85] |
| ENSG00000199683 | RN7SKP185 [miscRNA] | 0.0864 | -7.7758 [0.6542] | 1.1201e-70 [1.5208e-70] |
| ENSG00000188643 | S100A16 [protein coding] | 0.0855 | 10.5736 [1.9572] | 1.3122e-59 [2.2729e-59] |
| ENSG00000128218 | VPREB3 [protein coding] | 0.0841 | 9.2479 [0.4986] | 1.0546e-106 [1.8266e-106] |
| ENSG00000200087 | SNORA73B [snoRNA] | 0.0834 | -7.3992 [0.9972] | 2.1027e-44 [3.6420e-44] |
| ENSG00000136531 | SCN2A [protein coding] | 0.0802 | -8.1226 [0.0530] | 2.0875e-11 [3.6157e-11] |
| ENSG00000267206 | LCN6 [protein coding] | 0.0797 | 7.7130 [0.5306] | 1.4949e-40 [2.5893e-40] |
| ENSG00000263667 | Novel gene [lncRNA] | 0.0794 | 7.9739 [0.6103] | 4.2172e-54 [7.3005e-54] |
| ENSG00000125845 | BMP2 [protein coding] | 0.0792 | 8.1838 [0.1371] | 3.4885e-54 [6.0421e-54] |
| ENSG00000254535 | PABPC4L [protein coding] | 0.079 | -7.1982 [0.7380] | 1.3277e-08 [2.2996e-08] |
| ENSG00000150722 | PPP1R1C [protein coding] | 0.079 | -7.7929 [0.1759] | 5.8328e-23 [1.0103e-22] |
| ENSG00000149256 | TENM4 [protein coding] | 0.0783 | 8.6083 [0.1512] | 7.0158e-50 [1.2152e-49] |

| | | | | |
|---|---|---|---|---|
| ENSG00000253883 | IGHV3-19 [pseudogene] | 0.0777 | -8.7809 [0.8666] | 7.6242e-46 [1.3205e-45] |
| ENSG00000187621 | TCL6 [lncRNA] | 0.0772 | 8.4657 [0.0559] | 3.4849e-71 [6.0361e-71] |
| ENSG00000276778 | LINC02227 [lncRNA] | 0.0765 | 7.0439 [1.3625] | 1.2184e-38 [2.1025e-38] |
| ENSG00000161544 | CYGB [protein coding] | 0.0763 | 7.9310 [0.0436] | 2.4930e-70 [3.8479e-70] |
| ENSG00000170558 | CDH2 [protein coding] | 0.076 | -5.1614 [2.2585] | 1.1728e-08 [1.4038e-08] |
| ENSG00000128918 | ALDH1A2 [protein coding] | 0.0757 | -8.6717 [1.1190] | 1.7997e-07 [3.1172e-07] |
| ENSG00000251381 | LINC00958 [lncRNA] | 0.0757 | 8.2410 [0.3514] | 6.3176e-57 [1.0942e-56] |

Abbreviations: ALL, acute lymphoblastic leukemia; DEA, differential expression analysis; log2FC, log2 fold change; SD, standard deviation; FDR, false discovery rate.
