## Supplementary Table S5 for "Data-driven discovery of gene expression markers distinguishing pediatric acute lymphoblastic leukemia subtypes"

| Supplementary Table S5. Selected ENSEMBL gene IDs from random forest variable selection together with their external gene name and biotype. Random forest was applied on two predicted clusters as found using the unsupervised clustering framework *cola* (predicted clusters 1 and 4). The out-of-bag (OOB) error is of the final model selected following the feature selection procedure. The final model is selected as the one containing the least amount of features with an OOB error within 1 standard error of the minimum OOB error of all fitted random forests. | | |
|---|---|---|
| **Seed run** | **ENSEMBL gene ID (external gene name) [biotype]** | **OOB error** |
| 1 | ENSG00000154723 (ATP5PF) [protein coding], ENSG00000236863 (RPL23AP23) [processed pseudogene] | 0 |
| 2 | ENSG00000118181 (RPS25) [protein coding], ENSG00000169020 (ATP5ME) [protein coding] | 0 |
| 3 | ENSG00000125356 (NDUFA1) [protein coding], ENSG00000239398 (RN7SL342P) [misc RNA] | 0 |
| 4 | ENSG00000136149 (RPL13AP25) [processed pseudogene], ENSG00000142676 (RPL11) [protein coding] | 0 |
| 5 | ENSG00000171530 (TBCA) [protein coding], ENSG00000182004 (SNRPE) [protein coding] | 0 |
| 6 | ENSG00000226360 (RPL10AP6) [processed pseudogene], ENSG00000280392 (novel gene) | 0 |
| 7 | ENSG00000225071 (RPS26P58) [processed pseudogene], ENSG00000243964 (RPL23AP65) [processed pseudogene] | 0 |
| 8 | ENSG00000108961 (RANGRF) [protein coding], ENSG00000264063 (novel gene) [miRNA] | 0 |
| 9 | ENSG00000118181 (RPS25) [protein coding], ENSG00000228861 (RPL23AP12) [processed pseudogene] | 0 |
| 10 | ENSG00000146066 (HIGD2A) [protein coding], ENSG00000185641 (novel gene) [processed pseudogene] | 0 |
| Abbreviations: TEC, To be Experimentally Confirmed; OOB, out-of-bag. | | |
