## Supplementary Table S6 for "Data-driven discovery of gene expression markers distinguishing pediatric acute lymphoblastic leukemia subtypes"

| **Supplementary Table S6. Hazard ratios of the defined subset of 14 subtype-related gene expression markers together with 95% confidence intervals and *p*-values. Hazard ratios were found from a univariate Cox regression model with gene expression as the explanatory variable and survival data as the response variable performed on the independent Danish cohort of patients with acute lymphoblastic leukemia (ALL). Those markers with NA values did not comply with the proportional hazards assumption.** | | | |
|---|---|---|---|
| **Gene** | **Hazard ratio [95% CI]** | ***p*-value** | **FDR** |
| **ENSG00000118523** | 0.8035 [0.6486-0.9953] | 0.0452 | 0.1356 |
| **ENSG00000128218** | 0.8111 [0.6629-0.9926] | 0.0422 | 0.1356 |
| **ENSG00000164100** | 1.0045 [0.7945-1.2702] | 0.9698 | 0.9698 |
| **ENSG00000164330** | 0.8071 [0.6572-0.9911] | 0.0408 | 0.1356 |
| **ENSG00000199683** | 1.0915 [0.8760-1.3599] | 0.4353 | 0.5401 |
| **ENSG00000199831** | 1.0981 [0.8613-1.3910] | 0.4501 | 0.5401 |
| **ENSG00000200312** | 1.1101 [0.8540-1.4429] | 0.4351 | 0.5401 |
| **ENSG00000201901** | 1.1076 [0.9106-1.3471] | 0.3064 | 0.5401 |
| **ENSG00000202058** | 0.8621 [0.5236-1.4193] | 0.5595 | 0.6104 |
| **ENSG00000223806** | 0.7924 [0.6398-0.9814] | 0.0330 | 0.1356 |
| **ENSG00000227706** | 0.8206 [0.6686-1.0073] | 0.0587 | 0.1409 |
| **ENSG00000271394** | 1.2041 [0.8300-1.7469] | 0.3279 | 1.2041 |
| **ENSG00000200087** | NA | NA | NA |
| **ENSG00000200959** | NA | NA | NA |
| Abbreviations: ALL, acute lymphoblastic leukemia; CI, confidence interval; FDR, false discovery rate | | | |
