## Supplementary Text S1 for "Data-driven discovery of gene expression markers distinguishing pediatric acute lymphoblastic leukemia subtypes"

*Updated GC content and gene length annotations in TCGAbiolinks*

We updated the current geneInfoHT table in TCGAbiolinks containing GC content annotations and gene lengths using the R packages biomaRt and EDASeq. This table is used in the *TCGAbiolinks_Normalization* function when normalizing the data for GC content or gene length. We used the *useMart* function to connect to the ENSEMBL BioMart database and the *hsapiens_gene_ensembl* dataset. Afterwards, we used the *getBM* function to obtain ENSEMBL gene IDs from ENSEMBL. Finally, we used the *getGeneLengthAndGCContent* function to obtain gene lengths and GC contents of the obtained list of ENSEMBL gene IDs. See the issue on GitHub for more details: https://github.com/BioinformaticsFMRP/TCGAbiolinks/issues/492. The original code developed by us is reported in the GitHub repository associated with this publication for consultancy. These changes are implemented in TCGAbiolinks and here, we used TCGAbiolinks version 2.24.3.
Comparing the results from the original table geneInfoHT from TCGAbiolinks and the updated table containing these GC content annotations led to the loss of 33410 and 236 genes, respectively. Thus, we gain more information thanks to the new sources of annotations. Subsequently, we examined in detail the 236 genes for which we could not retrieve annotations in BioMart to ensure we were not losing any essential genes. These genes were all retired from the current ENSEMBL database, hence their lack of GC content annotation (see GitHub repository). The majority of these genes were long non-coding RNAs, pseudogenes, and only 53 protein coding genes (see **Supplementary Table S1** for information of these lost genes).
