## Supplementary Text S2 for "Data-driven discovery of gene expression markers distinguishing pediatric acute lymphoblastic leukemia subtypes"

*Pipeline settings of RNA sequencing pipeline*

The pipeline is using Snakemake workflow manager [1] ensuring high level of reproducibility, portability, and parallelization. The pipeline is available on GitHub: https://github.com/ELELAB/RNA_DE_pipeline.

*Trimming*
Reads were trimmed using Cutadapt [2]. Specifically, adapters and reads shorter than 35 bp were removed.
Params:
`"--minimum-length 35 -a AGAGCACACGTCTGAACTCCAGTCAC -g AGATCGGAAGAGCACACGT -A AGAGCACACGTCTGAACTCCAGTCAC -G AGATCGGAAGAGCACACGT"`

*Read mapping*
As a first step, reference genome hg38 release 104 was indexed using STAR [3] using transcript annotations GENCODE v38.
Params:
`"--sjdbGTFfile resources/genome.gtf --sjdbOverhang 149"`

Next, pair-end reads were aligned using STAR aligner onto the reference genome. Multimapping reads were retained in order to keep information about possible contamination e.g. by rRNA.
Params:
`"--twopassMode Basic --twopass1readsN -1 --sjdbOverhang 149 --sjdbGTFtagExonParentGene gene_name --outSAMtype BAM Unsorted --quantMode GeneCounts --outFilterMultimapNmax 200"`

*Alignment sorting*
Alignments were sorted by coordinate using Picard (*Picard Tools - By Broad Institute*).
Params:
`"VALIDATION_STRINGENCY=LENIENT CREATE_INDEX=true"`

*Calculating gene counts*
Gene counts were calculated using FeatureCounts from the SubRead package [4].
Params:
`"-p -t exon -g gene_id -s 2 -B -C --minOverlap 60"`

*Quality control*
Quality control (QC) is performed on raw fastq files using FastQC (*Babraham Bioinformatics - FastQC A Quality Control Tool for High Throughput Sequence Data*). QC based on read alignment was provided by Picard (CollectRnaSeqMetrics, CollectHsMetrics,

CollectAlignmentSummaryMetrics, CollectInsertSizeMetrics, CollectQualityDistributionMetrics, MeanQualityByCycle, CollectBaseDistributionByCycle, CollectGcBiasMetrics, CollectQualityYieldMetrics), RSeQC (Junction Annotation, Junction Saturation, Bam Stat, Read Distribution, Inner Distance, Read GC Content, Read Duplication, Infer Experiment) [5] and STAR. Extensive quality control results are wrapped into a single report using MultiQC [6].
